# ETS1, a target gene of the EWSR1::FLI1 fusion oncoprotein, regulates the expression of the focal adhesion protein TENSIN3

**DOI:** 10.1101/2023.12.21.572864

**Authors:** Vernon Justice Ebegboni, Tamara L. Jones, Tayvia Brownmiller, Patrick X. Zhao, Erica C. Pehrsson, Soumya Sundara Rajan, Natasha J. Caplen

**Affiliations:** Functional Genetics Section, Genetics Branch, Center for Cancer Research, National Cancer Institute, National Institutes of Health, Bethesda, MD 20892, USA; Omics Bioinformatics Facility, Genetics Branch, Center for Cancer Research, National Cancer Institute, National Institutes of Health, Bethesda, MD 20892, USA; Advanced Biomedical Computational Science, Frederick National Laboratory for Cancer Research, Frederick, MD 21702, USA

**Keywords:** EWSR1::FLI1, ETS1, Ewing sarcoma, Transcriptional repression, TENSIN3, TNS3

## Abstract

The mechanistic basis for the metastasis of Ewing sarcomas remains poorly understood, as these tumors harbor few mutations beyond the chromosomal translocation that initiates the disease. Instead, the epigenome of Ewing sarcoma (EWS) cells reflects the regulatory state of genes associated with the DNA binding activity of the fusion oncoproteins EWSR1::FLI1 or EWSR1::ERG. In this study, we examined the EWSR1::FLI1/ERG’s repression of transcription factor genes, concentrating on those that exhibit a broader range of expression in tumors than in EWS cell lines. Focusing on one of these target genes, *ETS1*, we detected EWSR1::FLI1 binding and an H3K27me3 repressive mark at this locus. Depletion of EWSR1::FLI1 results in ETS1’s binding of promoter regions, substantially altering the transcriptome of EWS cells, including the upregulation of the gene encoding TENSIN3 (TNS3), a focal adhesion protein. EWS cell lines expressing ETS1 (CRISPRa) exhibited increased TNS3 expression and enhanced movement compared to control cells. The cytoskeleton of control cells and ETS1-activated EWS cell lines also differed. Specifically, control cells exhibited a distributed vinculin signal and a network-like organization of F-actin. In contrast, ETS1-activated EWS cells showed an accumulation of vinculin and F-actin towards the plasma membrane. Interestingly, the phenotype of ETS1-activated EWS cell lines depleted of TNS3 resembled the phenotype of the control cells. Critically, these findings have clinical relevance as *TNS3* expression in EWS tumors positively correlates with that of *ETS1*.

**Significance:** ETS1’s transcriptional regulation of the gene encoding the focal adhesion protein TENSIN3 in Ewing sarcoma cells promotes cell movement, a critical step in the evolution of metastasis.

**Graphical abstract:** 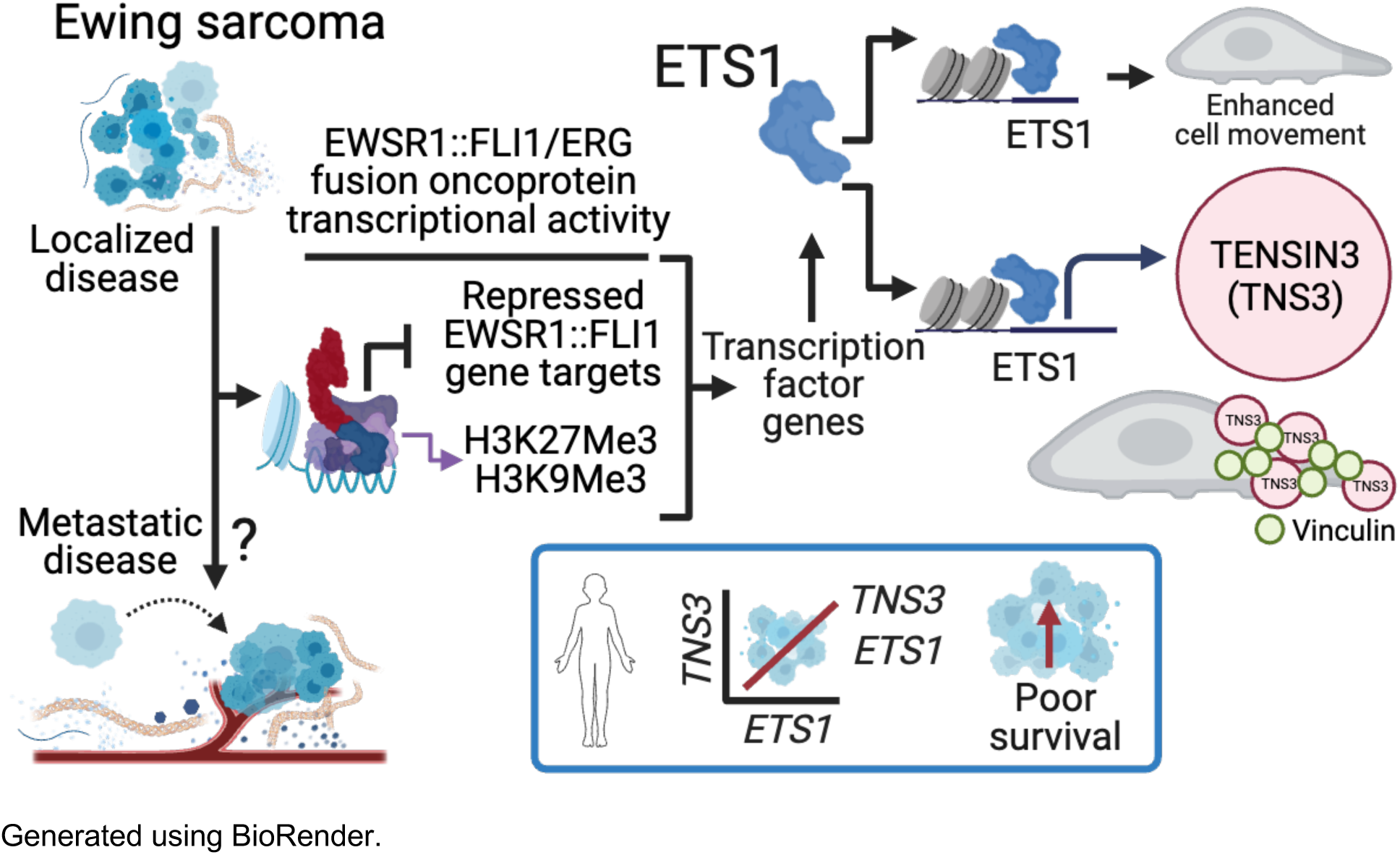

## Introduction

Since its discovery in the 1920s, the treatment of Ewing sarcoma (EWS), an aggressive bone and soft tissue sarcoma affecting children and young adults, has improved dramatically, and the current five-year survival rate for patients with localized disease is between 65 and 75% (1). However, patients with metastatic disease at diagnosis, typically pulmonary and osseous metastases, have five-year survival rates of about 50% and less than 30%, respectively (1). Highlighting the urgent need for the development of treatment strategies targeting EWS metastatic disease, an epidemiological study of over 800 EWS and EWS-like tumor cases identified that 35% of patients had detectable metastatic disease at presentation (2). However, the advancement of such therapies will require an enhanced understanding of the biological basis for the propensity of primary EWS tumor cells to disseminate and the processes that govern the generation of metastatic lesions.

In most solid malignancies, one determinant of the phenotypic plasticity required for metastasis is the genetic complexity of the primary tumor (3). In contrast, in EWS, a single oncogenic mutation drives tumor development. In most cases of EWS, the primary genetic event involves either a t(11:22) or t(21:22) translocation (4,5). These translocations result in the expression of either the EWSR1::FLI1 or EWSR1::ERG fusion oncoproteins that function as aberrant transcription factors due to the presence of the FLI1- or ERG-DNA binding domains of the ETS family of proteins (reviewed in 6). Importantly, Ewing sarcomas harbor few additional mutations, and thus, there is a need to determine the alternative mechanisms by which EWS cells attain the phenotypic features required for metastasis.

The EWSR1::FLI1/ERG fusion oncoproteins function as activators and repressors of gene expression (7). Several recent studies have suggested that genes typically repressed by EWSR1::FLI1/ERG fusion proteins may become activated due to variable fusion protein levels or external stimuli and that such changes in gene expression may contribute to the development of metastatic EWS (8–11). Repressed EWSR1::FLI1/ERG gene targets include many encoding proteins involved in cell-cell and cell-extracellular interactions, specifically proteins involved in actin cytoskeletal organization, extracellular matrix (ECM)-receptor interactions, and focal adhesion formation – processes associated with cell migration and invasion (9). Altered expression of repressed EWSR1::FLI1/ERG gene targets because of variable activity of the fusion protein within a tumor, effects of the microenvironment on tumor cell signaling, or other factors could, thus, result in the phenotypic changes associated with processes that promote the dissemination of tumor cells and consequently metastasis (reviewed in 12).

Current limitations to the study of EWS metastasis include the lack of statistically powered clinical datasets that match tumor samples from primary and metastases and the paucity of *in-vivo* models of EWS metastasis. In this study, we have harnessed alternative resources, specifically tractable EWS cell lines, and tumor gene expression profiles, to evaluate the hypothesis that repressed regulators of cell differentiation can contribute to EWS cell dissemination if activated, even transiently, and independent of the mechanism by which this occurs. Utilizing epigenomic and transcriptomic approaches, we have identified *ETS1* as a repressed gene target of EWSR1::FLI1. Using ectopic expression of ETS1 and CRISPR activation (CRISPRa) of the endogenous gene, we demonstrate that ETS1’s regulation of gene expression is distinct from that of EWSR1::FLI1 and that its expression induces substantial transcriptomic changes, including the increased expression of the focal adhesion associated protein TENSIN3 (TNS3) and enhanced cell movement and migration. Critically, using multiple datasets, we observe a positive correlation between *ETS1* and *TNS3* RNA levels in EWS tumor samples, suggesting that EWS tumor cells expressing ETS1 have the potential to exhibit a phenotype that promotes cell movement.

## Materials and Methods

### Cell lines and reagents

**Supplemental Table S1** details the cell lines, siRNAs and PCR primers, plasmids, and other critical reagents used in this study. Cells were cultured in RPM1-1640 or DMEM media (Thermo Fisher Scientific, Waltham, MA) supplemented with 10% FBS and Plasmocin Prophylactic (Invivogen, San Diego, CA) and grown at 37°C, 5% CO_2_. We confirmed the identity of cell lines using short tandem repeat (STR) analysis (ATCC; details available upon request), and we monitored for mycoplasma contamination using the MycoAlert Plus system (Lonza, Walkersville, MD).

### RNAi, qRT-PCR analysis, and RNA sequencing

Cells were reverse transfected, and total RNA was isolated and cDNA synthesized as previously described (13–15). Quantitative Real-Time PCR (qRT-PCR) was performed using PowerUp SYBR Green Master Mix (A25778; Thermo Fisher Scientific) on an ABI StepOne Plus Real-Time PCR system (Applied Biosystems, Foster City, CA) or a CFX384 (Bio-Rad, Hercules, CA). Fold-change gene expression was calculated by the ^ΔΔ^CT method and normalized to *NACA* mRNA levels. Samples were generated from three independent experiments unless stated otherwise. For paired-end RNA sequencing (RNA-seq), RNA was extracted (Maxwell 16 LEV simplyRNA purification kit, Promega, *Madison, WI*), and libraries were prepared and sequenced using standard protocols and a Nextseq 2000 instrument (Illumina, San Diego, CA). The CCR Collaborative Bioinformatics Resource (CCBR) RNA-seq pipeline was used for data analysis https://bioinformatics.ccr.cancer.gov/ccbr/pipelines-software/ccbr-pipeliner/ (see **Supplemental Materials and Methods** for additional details).

### Generation of plasmids and stable cell lines

To generate the *ETS1*-pLOC construct, a synthetic human *ETS1* cDNA (NM_001143820) with custom flanking restriction enzyme sites (GENEWIZ, South Plainfield, NJ) was cloned into the pLOC vector (OHS5832; Dharmacon/Horizon Discovery, Cambridge, UK). Lentivirus was produced in HEK-293T cells using either the Trans-Lentiviral ORF Packaging Kit (Dharmacon/Horizon Discovery) or Lipofectamine 3000 (Thermo Fisher Scientific). Viral supernatants were collected 72 hours (hrs) post-transfection, concentrated (PEG Virus Precipitation Kit, Abcam, *Waltham, MA*), and added dropwise to cells in the presence of media supplemented with 8 µg/ml polybrene. Transduced cells were selected using puromycin (2 µg/ml, Thermo Fisher Scientific). For the generation of CRISPR lines, cells were transduced with virus expressing dCas9-VP64 (Addgene, Watertown, MA), selected using blasticidin (6 – 10 µg/ml, Thermo Fisher Scientific), and subjected to single-cell sorting. Plasmids expressing *ETS1* sgRNAs (see **Supplemental Table S1**) were co-electroporated into the dCas9-VP64-expressing cells using the Amaxa^TM^ Cell Line Nucleofector™ Kit R (Lonza, VCA-1001) and further subjected to puromycin selection and single-cell cloning. CRISPR-activation efficiency was assessed by qPCR and/or immunoblotting.

### Chromatin Immunoprecipitation (ChIP) and CUT&RUN analysis

ChIP assays were performed using the SimpleChIP plus Enzymatic Chromatin IP kit (9005S; Cell Signaling Technology, Danvers, MA) following the manufacturer’s protocol. For quantitative ChIP-PCR-based analysis, the ChIP-enriched DNA was analyzed by using the primers listed in **Supplemental Table S1**. For ChIP-sequencing (ChIP-seq) and Cleavage Under Targets and Release Using Nuclease (CUT&RUN), two independent replicates of each sample were prepared except for the ChIP-seq analysis of H3K27ac, which was prepared in triplicate. See **Supplemental Materials and Methods** for detailed descriptions of the ChIP-seq and CUT&RUN protocols (the latter carried out as previously described (16)) and the analytical pipelines used to examine these datasets (https://bioinformatics.ccr.cancer.gov/ccbr/pipelines-software/ccbr-pipeliner/; https://github.com/CCBR/CARLISLE).

### GGAA motif quantification

Quantification of the GGAA composition of EWSR1::FLI1 and ETS1 peaks was performed as previously described (17) with modifications. Briefly, the genomic sequence within a range of 500 bp around the summit of EWSR1::FLI1 or ETS1 peaks called by MACS2 was extracted (hg38). The number of (GGAA)_n_ or (TTCC)_n_ repeats (from 1-4 or >4 consecutive GGAA motifs without any gap) in each 500 bp range counted.

### Immunoblotting and Immunofluorescence

Whole-cell lysates were prepared using cell extraction buffer (62 mM Tris-HCl, pH 8.0, 2% SDS, 10% Glycerol) and sonicated. Protein concentrations were determined using a BCA assay (Thermo Fisher Scientific). 20 – 30 µg of each protein sample was analyzed following standard immunoblotting protocols (see **Supplemental Materials and Methods**) using antibodies at the dilutions detailed in **Supplemental Table S1**. For analysis using immunofluorescence (IF), SK-N-MC and ES-5838 (2 x 10^5^) cells were plated in each well of a 6-well plate on round coverslips and grown for 72 to 96 hrs. Cells grown on the coverslips were processed using standard procedures (see **Supplemental Materials and Methods**) and analyzed using antibodies at the dilutions detailed in **Supplemental Table S1** and the DAPI stain to define nuclei.

### Cell proliferation and trans-well chemotaxis migration assays

For proliferation assays, SK-N-MC and ES5838 (4 x 10^4^) cells per well were plated in a 24-well plate and placed in an IncucyteS3 incubator (Sartorius, Essen BioScience, Inc. Ann Arbor, MI). Nine images per well were taken every 6 hrs for seven days. Images were analyzed using the Incucyte 2020B GUI Analysis Software. For trans-well chemotaxis migration assays, 2.5 x 10^3^ cells/well were plated in FBS-free media in the upper chamber of a 96-well Incucyte ClearView cell migration plate (8 µm pore size; #4582 Sartorius). Cells were incubated at 37°C for 45 minutes to allow cells to settle on the membrane before adding RPMI-1640 media supplemented with 10% FBS to the bottom chamber. The cells were placed in an IncucyteS3 incubator at 37°C with images of top/bottom chambers taken every 6 hrs for at least 72 hrs. The total area of cells at the bottom chamber was normalized to the initial top chamber area to quantify the migratory phenotype.

### Cell motility assay and tracking analysis

SK-N-MC and ES-5838 (1.8 x 10^3^ cells/well) were plated in 96 well ImageLock microplate (#4379 Sartorius). Cells were incubated at 37°C overnight to allow cells to settle, placed in an IncucyteS3 incubator (Sartorius), and images were taken every 2 hrs for 100 hrs. Two phase contrast images per well were captured at each time point at 20X magnification. Images up to 96 hrs across representative wells were compiled into a stack in Fiji (18). For each stack, an ROI of 800 x 800 pixels was defined, and nuclei within this ROI were segmented using Otsu thresholds in Fiji. The distance traveled was measured using the TrackMate plugin in Fiji. Only nuclei with a mean quality >100 across all time points with a minimum duration of ten occurrences in 96 hrs were assessed. Distance traveled per segmented nuclei over 96 hrs in each representative well was obtained from TrackMate analysis, exported to Prism, and plotted as shown. Phase contrast movies (two frames per second) were generated using Fiji. Confluency data from all the wells from each sample was analyzed as previously described.

### External data sets

Normalized RNA-seq data (TMM-RPKM - trimmed mean of M values-reads per kb per million mapped reads) for 79 EWS tumors and 42 EWS cell lines were supplied by Dr Javed Khan, Genetics Branch, CCR. These data are deposited in dbGAP submissions phs001928.v1.p1, phs000768, and phs001052, and combined in https://oncogenomics.ccr.cancer.gov/production/public/viewProjectDetails/24421, and described in Brohl *et al*., 2021 (19). **Supplemental Table S1** includes a list of the 42 EWS cell lines within this dataset. The R2 Genomics Analysis and Visualization Platform (http://r2.amc.nl) (20) was used to evaluate deposited EWS tumor expression profiles assessed using microarray-based platforms. Data reported from the following cohorts were used: “Gene Expression Profiling of Ewing Sarcoma Tumors Reveals the Prognostic Importance of Tumor-Stromal Interactions: A Report from the Children’s Oncology Group” GSE63157 (85 tumors) (21); “ Expression profiling of Ewing sarcoma samples” GSE34620 (117 tumors) (22); “Overcoming resistance to conventional drugs in Ewing’s sarcoma and identification of molecular predictors of outcome” GSE12102 (37 tumors) (23); “Expression profiling of Ewing sarcoma samples” GSE142162 79 samples (24).

### Statistical and graphical analyses

Calculations were performed in Excel (Microsoft), and data were exported to Prism 8.0.0 for Mac (GraphPad, La Jolla, CA) for statistical analyses. Unless stated otherwise, results are shown as mean ± standard error of the mean (SEM). A p-value of <0.05 was considered significant, though, for some analyses, more stringent criteria were applied. Standard R functions and packages (ggplot2, dplyr, ggrepel, and VennDiagram) were used to perform other analyses and to generate plots.

### Data availability

The data and resources generated are available upon reasonable request by contacting the corresponding author. The RNA-seq, ChIP-seq, and CUT&RUN datasets reported here are available in the Gene Expression Omnibus at GSE243184.

## Results

### EWSR1::FLI1 represses the expression of multiple transcriptional factor genes

To identify transcription factor genes repressed by the EWSR1::FLI1 fusion oncoprotein, we silenced the fusion oncogene in TC-32, TC-71, and A673 EWS cell lines (**Supplemental Fig. S1A**) and identified the differentially expressed genes (DEGs) (±1.5-fold change; FDR <0.05) (**Supplemental Fig. S1B, Supplemental Table S2**). The DEGs observed in all three EWS cell lines showed enrichment for genes regulating the organization of the extracellular matrix, cell adhesion, and migration as showing an increased expression following EWSR1::FLI1 depletion (1107 genes) and enrichment for genes that function in the cell cycle, showing decreased expression (1111 genes) (**Supplemental Fig. S1C, Supplemental Table S2**). Using the dataset of 2218 genes regulated by EWSR1::FLI1, we examined a curated list of genes encoding proteins with *bona-fide* DNA binding activity (25) and determined that 128 encode transcription factors (**Supplemental Table S3**). Of these 128 genes, 61 exhibited decreased expression, including *HOXD13*, *NKX2-2*, *NR0B1*, and *SOX2*, previously linked to Ewing sarcoma biology (**Supplemental Fig. S1D**) (26–28). Of the upregulated genes after EWSR1::FLI1 silencing, 67 encoded transcription factors, including established regulators of cell differentiation and developmental processes, such as *SNAI2, ETS1,* and *RUNX2* (**Fig. 1A**).

**Figure 1:**
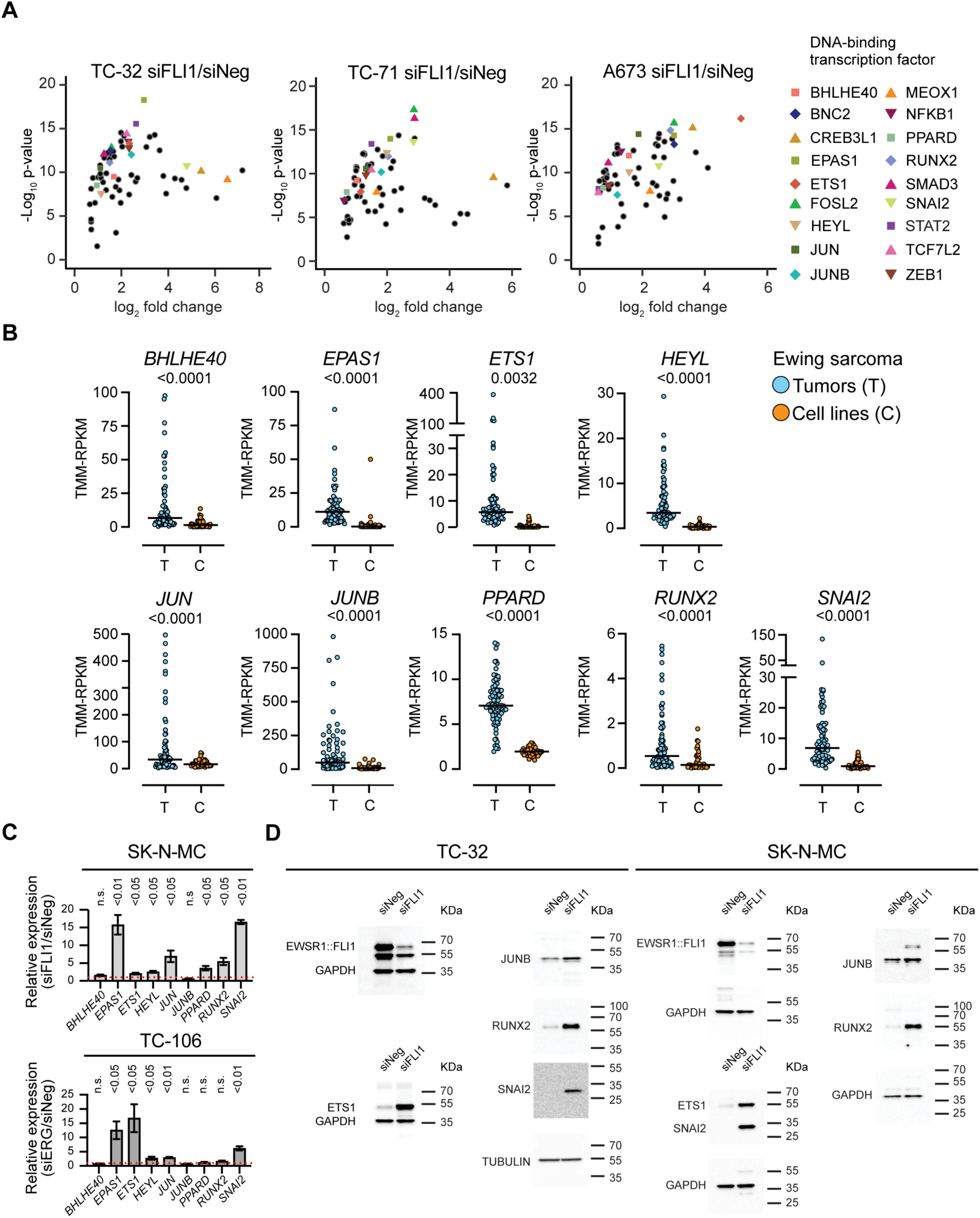
The EWSR1::FLI1/ERG fusion oncoproteins repress multiple transcription factor genes in EWS cell lines. (**A**) Differential expression of transcription factor genes that demonstrated a significant increase in expression following the silencing of *EWSR1::FLI1* in TC-32, TC-71, and A673 EWS cells (three biological replicates; fold change >1.5, FDR < 0.05). (**B**) Comparative expression of selected transcription factor genes in EWS tumor samples (n=79, blue circles) and EWS cell lines (n=42, orange circles) (19) (p-values determined using an unpaired t-test with Welch’s correction). (**C**) qRT-PCR analysis of a subset of EWSR1::FLI1-repressed genes in SK-N-MC (siFLI1-transfected) and TC-106 (siERG-transfected) EWS cells, relative to control (siNeg-transfected) (mean ± SEM from three independent experiments; p-values determined using multiple unpaired tests). (**D**) Immunoblots of whole-cell lysates prepared from the indicated cell lines transfected with the indicated siRNA (48 h) and analyzed using the antibodies against the indicated proteins.

To assess the potential clinical relevance of EWSR1::FLI1’s regulation of the transcription factor genes that exhibit an increase in expression following the depletion of the fusion oncoprotein, we examined normalized RNA-seq data from EWS tumors (n=79) and cell lines (n=42) (19). The distribution of gene expression in tumors relative to cell lines was higher (p<0.01) for 37 of the 67 transcription factor genes that exhibited an increase in expression following the silencing of *EWSR1::FLI1* in EWS cell lines. **Fig. 1B** summarizes the results for nine genes encoding different classes of transcription factors: the basic helix-loop-helix proteins BHLHE40, EPAS1, and HEYL; the bZIP-proteins JUN and JUNB; the nuclear receptor C4-type zinc finger, Runt-domain containing, and Cys2–His2 zinc finger proteins, PPARD, RUNX2, and SNAI2, respectively; and, like FLI1 and ERG, a member of the ETS family of proteins, ETS1. **Supplemental Fig. S2A** summarizes the results for the other 28 transcription factor genes. Based on these results, such genes may exhibit more variable expression in EWS tumors than cell line data suggest. To gain additional evidence that EWSR1::FLI1 represses the expression of specific transcription factors and extend our analysis to the EWSR1::ERG fusion protein, we examined the expression of the nine genes highlighted in **Fig. 1B** in two additional EWS cell lines, observing a significant increase in the expression of seven genes in SK-N-MC cells following silencing of *EWSR1::FLI1* and five genes in TC-106 cells post-silencing of *EWSR1::ERG*, including *ETS1* and *SNAI2*. (**Fig. 1C**, **Supplemental Fig. S2B**). Analysis of the proteins encoded by four of these genes, ETS1, SNAI2, JUNB, and RUNX2, confirmed that depletion of *EWSR1::FLI1* in TC-32 and SK-N-MC cells increases their expression (**Fig. 1D**). These results, emphasize EWSR1::FLI1/ERG’s repression of multiple transcription factors but also suggest that the degree of repression, as well as the resulting phenotypic effects, may vary in tumors.

### EWSR1::FLI1 represses the expression of *ETS1* and other regulators of cell differentiation

To investigate EWSR1::FLI1’s repression of specific genes, including those encoding transcription factors, we assessed its associations with distinct epigenetic states (**Supplemental Table S4).** Using samples generated from TC-32 cells, we employed CUT&RUN to determine baseline EWSR1::FLI1 binding sites (two independent experiments, two biological replicates each) indicated as A and B, respectively – **Figs. 2A** and **2B**) and mapped histone modifications associated with transcriptional activation, H3K27ac (**Fig. 2A),** and repression, H3K9me3 and H3K27me3 **(Fig. 2B)**. Complementing these analyses, we conducted ChIP-seq analyses of H3K27ac in the presence and absence of EWSR1::FLI1 (**Supplemental Fig. S3A**). We identified over 25,000 EWSR1::FLI1 binding sites, which overall exhibited enrichment for the ETS consensus sequence motif (**Supplemental Fig. S3B**) and the occupancy of multimeric GGAA sequences (≥ 4 GGAA repeats) previously reported for the fusion oncoprotein (**Fig. 2C**) (29). Also consistent with published results (7,30), we observed the predominate binding of EWSR1::FLI1 to genomic sites present in distal regions of genes and their H3K27ac peaks (**Supplemental Fig. S3C**). Examining EWSR1::FLI1 binding and epigenetic marks in the vicinity of the well-characterized EWSR1::FLI1 activated gene *CCND1*, we observed EWSR1::FLI1’s binding of a site 5’ of the locus, similar distributions of H3K27ac detected by CUT&RUN and ChIP-seq, and a decrease in the latter following depletion of EWSR1::FLI1 (**Supplemental Fig. S3D**). Next, we examined the distribution of the H3K9me3 and H3K27me3 modifications and mapped over 30,000 genomic regions associated with H3K9me3 marks and over 20,000 with H3K27me3. As an example, we examined the epigenetic regulation of *PHLDA1*, a gene bound by EWSR1::FLI1 that shows a substantial increase in gene activation as indicated by the H3K27ac mark in the absence of EWSR1:FLI1 (**Supplemental Fig. S3E)**. In the case of *PHLDA1,* we observed H3K27me3 across the gene body, suggesting that at this locus, EWSR1::FLI1 binding results in the recruitment of polycomb-repressive marks.

**Figure 2:**
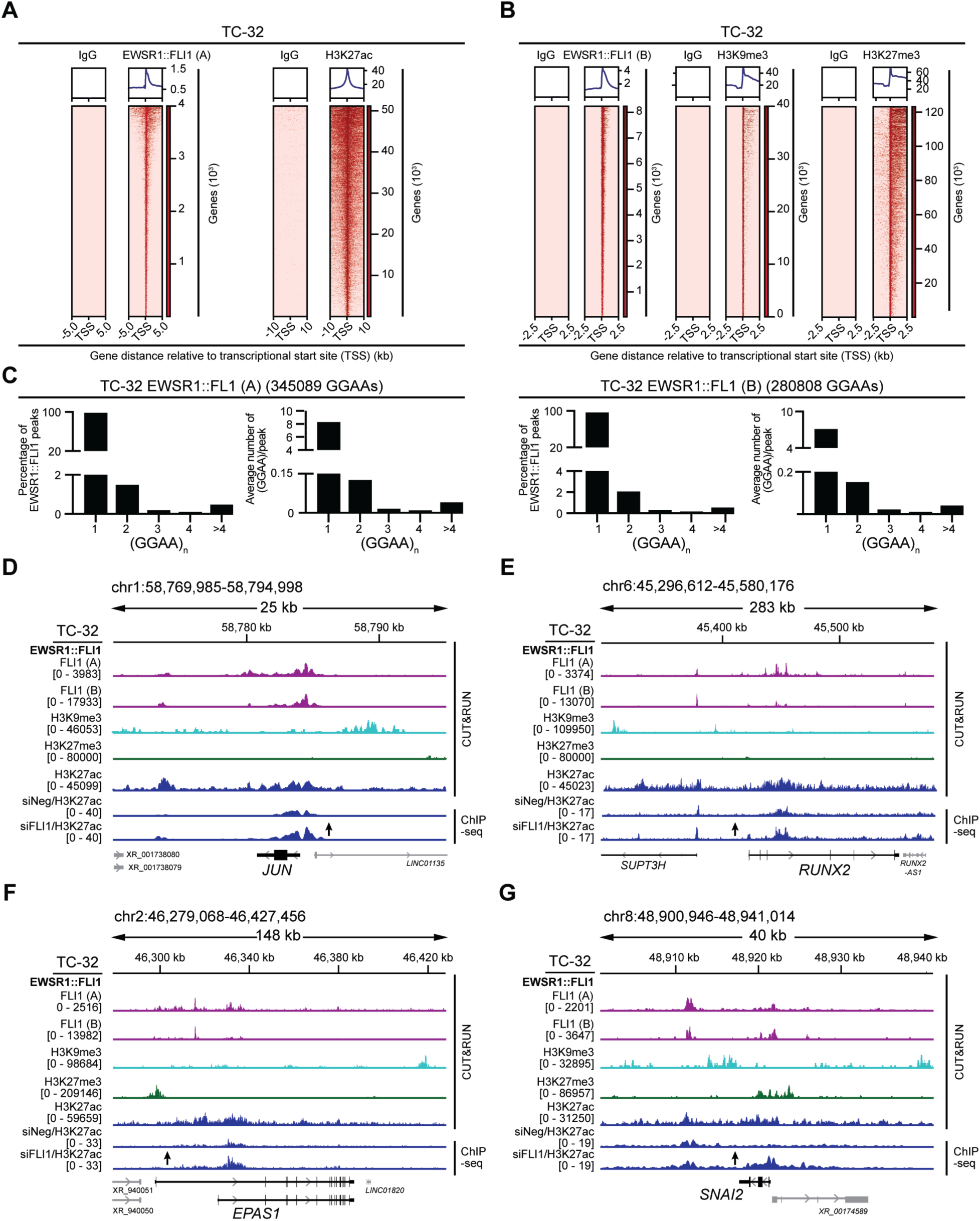
EWSR1::FLI1’s repression of transcription factor genes. (**A**) Genome-wide heatmaps of EWSR1::FLI1 (experiment A; n=2) and H3K27ac CUT&RUN-seq peak-centered signals in TC-32 cells showing 5kb and 10kb windows, respectively. (**B**) Genome-wide heatmaps of EWSR1::FLI1 (experiment B; n=2), H3K9me3, and H3K27me3 CUT&RUN-seq peak-centered signals in TC-32 cells showing 2.5kb windows. (**C**) The percentage of peaks with at least one (GGAA)_n_ (left) and the average number of (GGAA)_n_ per peak (right) in EWSR1::FLI1 binding sites based on consecutive GGAA repeat sequence motifs of 1, 2, 3, 4, or over 4. (**D-G**) EWSR1::FLI1 binding, and the H3K9me3, H3K27me3, and H3K27ac CUT&RUN-seq signals detected in unmodified TC-32 cells, and the H3K27ac ChIP-seq signals in control and siFLI1-transfected TC-32 cells at (**D**) the *JUN* locus, (**E**) the *RUNX2* locus, (**F**) the *EPAS1* locus, and (**G**) the *SNAI2* locus.

Focusing on the transcription factor genes with upregulated expression following depletion of EWSR1::FLI1 (**Fig. 1A**), we interrogated the binding of EWSR1::FLI1, H3K9me3, and H3K27me3 modifications at 64 of these loci (**Supplemental Table S5**). Using standard annotations (MACS2 narrow peaks, p-value <0.01 in both CUT&RUN datasets), 60 of the 64 loci showed evidence of EWSR1::FLI1 binding in the proximity of the gene body, albeit one of which exhibited a range of expression in the EWS tumor samples when compared to cell lines. Next, we examined the annotations of repressive marks (MACS2 broad peaks, p-value <0.01), which assigned H3K9me3 at 23 of the 60 transcription factor genes (∼38%), H3K27me3 at nine genes (∼15%), and both H3K9me3 and H3K27me3 in the proximity of seven genes (∼12%). The remaining loci showed no annotated marks. As examples, the *JUN* and *RUNX2* loci (**Figs. 2D, 2E**) show evidence of H3K9me3 modifications in their proximity. In contrast, the *EPAS1* and *SNAI2* loci (**Figs. 2F, 2G**) showed evidence of H3K9me3 and H3K27me3 modifications, though in both cases, examination of the relative location of these marks showed H3K27me3 as the closest peak. At these loci, we observed similar distributions of the H3K27ac signals whether detected by CUT&RUN or ChIP-seq; however, due to the technical differences between these methods, the magnitude of their signals differed substantially. Nevertheless, and consistent with our analysis of their gene expression following the depletion of EWSR1::FLI1, we observed a substantial increase in the H3K27ac signals present at each locus following the silencing of *EWSR1::FLI1*.

To obtain additional evidence that alterations in the transcriptional regulators our studies delineated as repressed by EWSR1::FLI1 are relevant to disease progression, we interrogated one of the few EWS tumor gene expression datasets (21) associated with disease outcome using the R2 Genomics Visualization platform (20) (**Supplemental Table S5)**. Intriguingly, this analysis showed high *ETS1* expression associated with poor overall survival probability (**Supplemental Table S5, Supplemental Fig. S4A**). Based on this finding, we chose to pursue *ETS1* as a candidate repressed target gene for further analyses, beginning with assessing its epigenetic regulation.

*ETS1* is located adjacent to *FLI1* on chromosome 11, and the alignment of RNA-seq data from EWSR1::FLI1 expressing EWS cell lines illustrates a striking juxtaposition of the effects of silencing the fusion oncogene and the expression of *ETS1*. As expected, we detect a decrease in the reads mapping to the 3’ *FLI1* exons following targeting of *EWSR1::FLI1*, whereas there is an increase in *ETS1* expression (**Fig. 3A**). Using TC-32-derived samples, we observed EWSR1::FLI1’s binding of multiple sites within *ETS1* **(Fig. 3B**), an observation we confirmed and extended to SK-N-MC-derived samples using ChIP-qPCR (**Fig. 3C**, **Supplemental Fig. S4B**).

**Figure 3:**
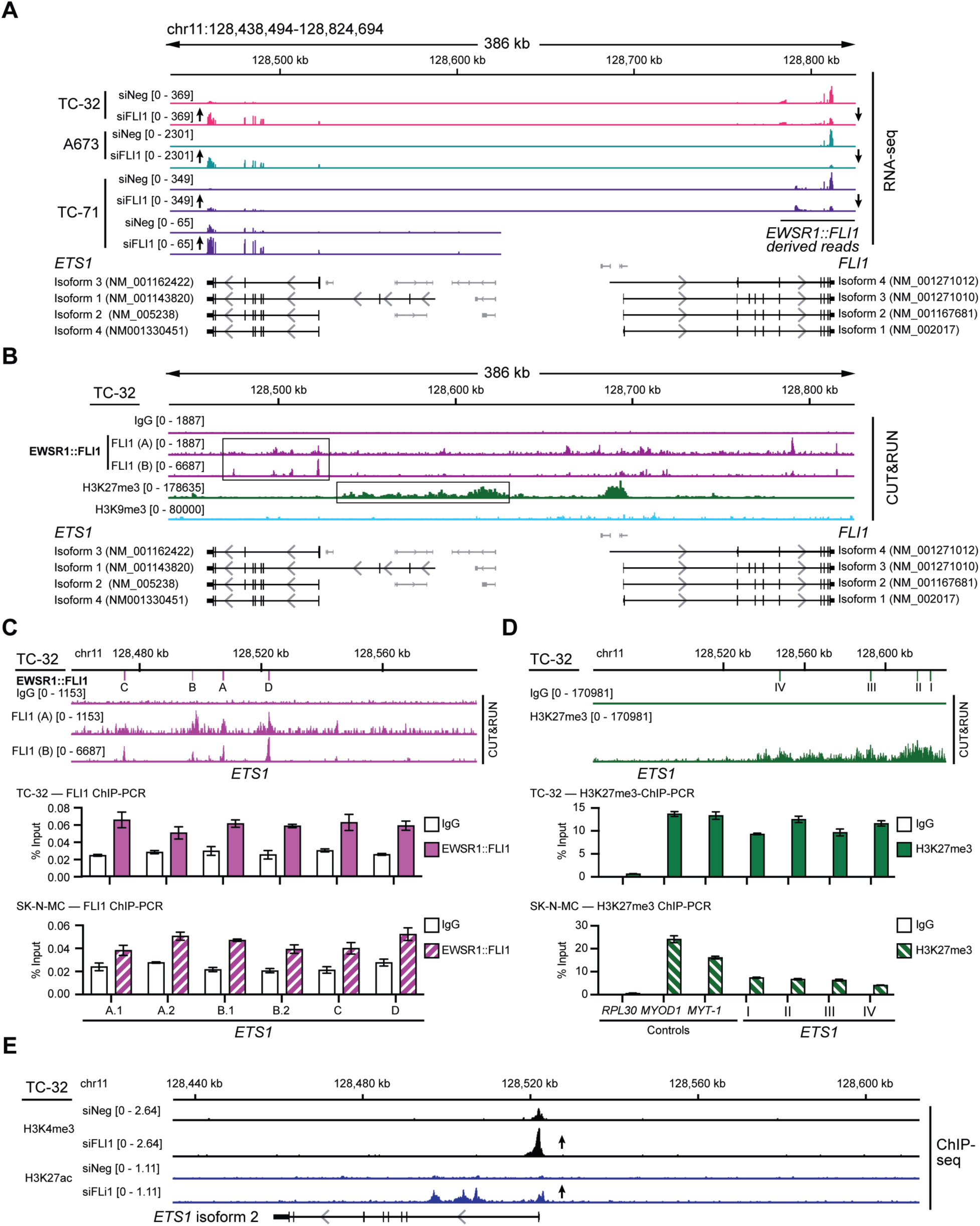
*ETS1* is a repressed gene target of EWSR1::FLI1. (**A**) RNA-seq read depth of control and *EWSR1::FLI1*-silenced TC-32, A673, and TC-71 cells focusing on the *ETS1* and *FLI1* loci. For clarity, two different scales show the read depth at the *ETS1* locus in TC-71 cells. (**B**) EWSR1::FLI1 binding, H3K27me3, and H3K9me3 modifications at the *ETS1* and *FLI1* loci of TC-32 cells. (**C**) ChIP-qPCR analysis of EWSR1::FLI1 binding at the indicated locations of the *ETS1* locus in TC-32 and SK-N-MC cells. See **Supplemental Fig. S4B** for sequence information. (**D**) ChIP-qPCR analysis of H3K27me3 deposition at the indicated locations of the *ETS1* locus in TC-32 and SK-N-MC cells. See **Supplemental Fig. S4C** for sequence information. (**E**) The *ETS1* locus showing H3K4me3 and H3K27ac modifications detected in control (siNeg) and siFLI1-transfected TC-32 cells.

Analysis of the H3K27me3 and H3K9me3 marks showed a distribution of H3K27me3 at a region adjacent to *ETS1* but not the latter mark (**Fig. 3B**). H3K27me3 ChIP-qPCR validated this finding and showed evidence for the same mark upstream of the *ETS1* locus in SK-N-MC cells (**Fig. 3D**, **Supplemental Fig. S4C**). Furthermore, we detected an increase in the H3K4me3 mark (**Supplemental Table S6, Supplemental Fig. S4D**) at the *ETS1* TSS and H3K27ac following silencing of the fusion oncogene (**Fig. 3E**), confirming transcriptional activity at this locus in the absence of EWSR1::FLI1. These analyses reveal that EWSR1::FLI1 directly represses another ETS family member, ETS1. Based on these results, we hypothesized that variabilities in EWSR1::FLI1 transcriptional activity may explain, at least in some cases, the range of *ETS1* mRNA levels observed in EWS tumors (**Fig. 1B; Supplemental Fig. S4A**). To investigate the consequences of ETS1 expression in EWS cells, we next assessed its binding in EWS cells depleted of EWSR1::FLI1.

### ETS1’s regulation of gene expression is distinct from that of EWSR1::FLI1

Following the depletion of EWSR1::FLI1, ChIP-seq analysis defined 2635 ETS1 binding sites in TC-32 cells (**Fig. 4A, Supplemental Table S7**). In contrast to EWSR1::FLI1, global ETS1 chromatin occupancy in EWSR1::FLI1-depleted cells predominantly mapped to promoter regions (∼84%) **(Fig. 4B)**. While ETS1 bound regions displayed enrichment for the canonical ETS DNA binding motifs **(Fig. 4C)**, ETS’s binding at GGAA motifs was predominately at single and interspersed GGAAs **(Fig. 4D)**. This is in contrast with EWSR1::FLI1’s binding at GGAA microsatellites and its aberrant function (7,31,32), and suggests that ETS1 genomic occupancy is distinct from that of EWSR1::FLI1.

**Figure 4:**
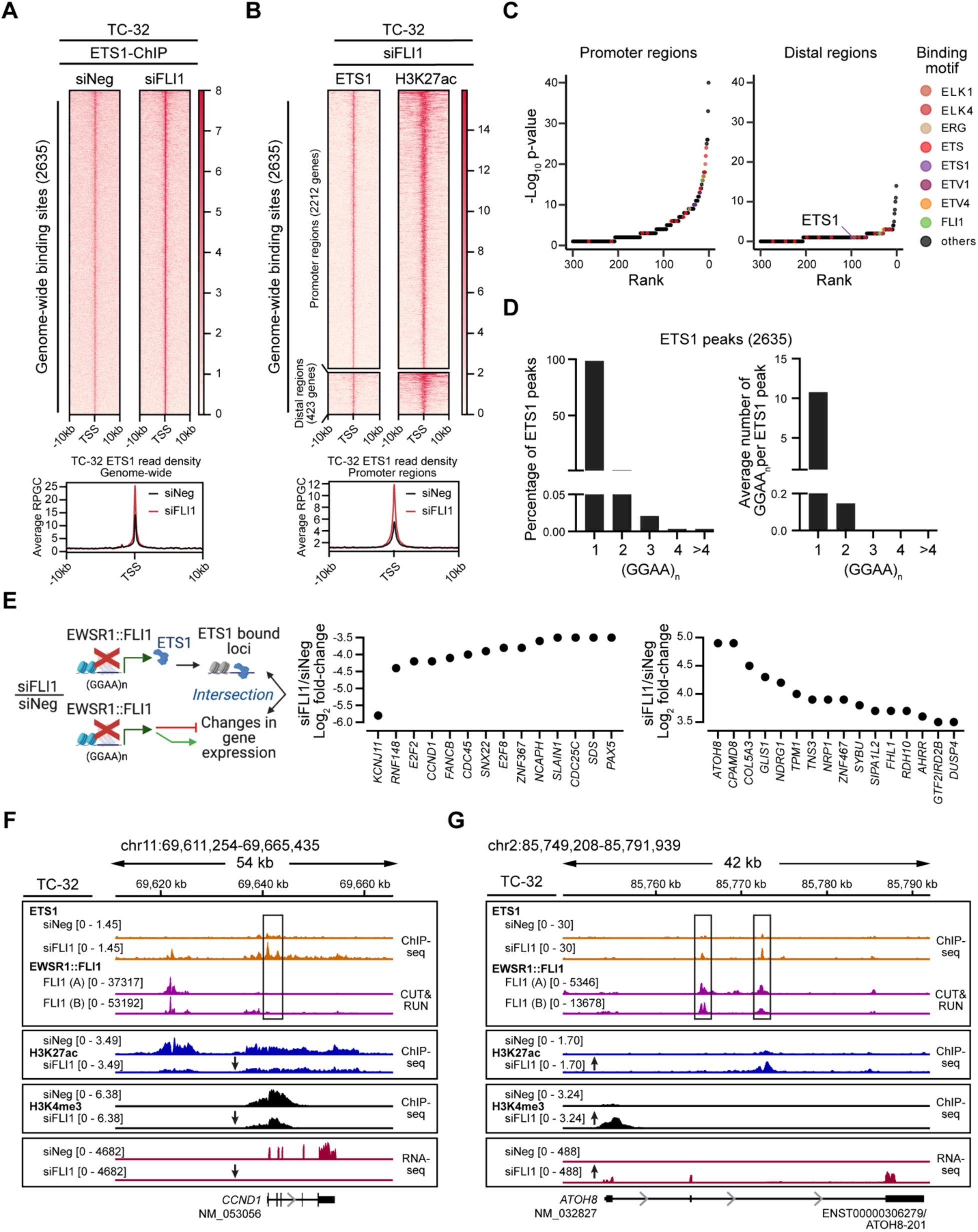
ETS1 DNA binding is distinct from that of EWSR1::FLI1. (**A**) Genome-wide heatmaps of ETS1 ChIP-seq peak-centered signals and read densities in control (siNeg) and *EWSR1::FLI1*-silenced (siFLI1) TC-32 cells showing 10kb windows. Rankings of the bound regions are based on the siFLI1 signal. (**B)** Cluster heatmaps of ChIP-seq peak-centered signals for ETS1 and H3K27ac in TC-32 siFLI1-transfected cells showing 10kb windows. Rankings of the cluster regions are based on the ETS1 signals, with the gene promoter regions defined as −3kb and +1kb from TSS and the distal regions as sites existing outside of promoter regions. Below the heatmap is a summary of the ETS1 read densities at promoter regions in control (siNeg) and *EWSR1::FLI1*-silenced (siFLI1) TC-32 cells. (**C**) Sequence motifs enriched in ETS1/H3K27ac promoter and ETS1/H3K27ac distal regions as defined in (**B**). (**D**) The percentage of peaks with at least one (GGAA)_n_ (left) and the average number of (GGAA)_n_ per peak (right) in ETS1 binding sites based on consecutive GGAA repeat sequence motifs of 1, 2, 3, 4, or over 4. (**E**) Schematic illustrating the intersectional analysis performed (left) and the changes in expression of ETS1-bound genes that are protein-encoding and exhibit changes in expression following depletion of EWSR1::FLI1 (greater or less than ±3.5 Log_2_-fold) (right). (**F)** The *CCND1* locus of TC-32 cells assayed under the indicated conditions showing ETS1 and EWSR1::FLI1 binding, H3K9me3, H3K27me3, H3K27ac, and H3K4me3 modifications, and gene expression. **(G)** The *ATOH8* locus of TC-32 cells assayed under the indicated conditions showing ETS1 and EWSR1::FLI1 binding, H3K9me3, H3K27me3, H3K27ac, and H3K4me3 modifications, and gene expression. The schematic in **E** was generated using BioRender.

To identify putative downstream targets regulated by ETS1 in the absence of EWSR1::FLI1, we intersected ETS1 occupancy and significantly deregulated genes from the RNA-Seq analysis of EWSR1::FLI1-depleted TC-32 cells **(Supplemental Fig. 1B)**. This analysis defined 958 genes as bound by ETS1, including the genes shown in **Fig. 4E**, which illustrates ETS1’s association with genes that exhibit decreased or increased expression following the silencing of *EWSR1::FLI1*. As an example of ETS1’s association with an activated EWSR1::FLI1 gene target, we examined its binding at the *CCND1* locus (**Fig. 4F**). Many studies have noted the high expression of CCND1, a critical regulator of the cell cycle, in EWS cells and its down-regulation following depletion of EWSR1::FLI1 (33–35). Our analysis confirmed the decrease in *CCND1* RNA levels in *EWSR1::FLI1*-silenced TC-32 cells, and a reduction in the H3K4me3 and H3K27ac marks at this locus. Interestingly, under control conditions, we detected EWSR1::FLI1 binding upstream of *CCND1* with no evidence of ETS1 binding. However, following the depletion of EWSR1::FLI1, we observed a less pronounced ETS1 peak in the same vicinity as EWSR1::FLI1 and a prominent ETS1 peak at the *CCND1* promoter. ETS1, like other ETS proteins, can repress and activate gene expression, leading us to speculate that at the *CCND1* locus, ETS1 may function as a transcriptional repressor. In contrast, ETS1 may promote the expression of *ATOH8* (**Fig. 4G**), which encodes a transcription factor recently implicated in the regulation of chondrogenic differentiation (36), cellular plasticity (37), and the differentiation and maintenance of skeletal muscle (38). Gene expression analysis and H3K4me3 and H3K27ac epigenetic marks all indicate that EWSR1::FLI1 suppresses the expression of *ATOH8* as each mark increases following depletion of the fusion oncoprotein. Consistent with ETS1 contributing to the promotion of *ATOH8* expression, we observed ETS1 binding at two regions within this locus. However, in this case these peaks coincided with EWSR1::FLI1 peaks, suggesting that ETS1 and EWSR1::FLI1 proteins may compete for binding at some sites.

### ETS1 regulates gene targets repressed by EWSR1::FLI1

We next interrogated ETS1’s regulation of gene expression in the presence of EWSR1::FLI1 by ectopically expressing ETS1 in TC-32 cells. We generated two single-cell clones, one that expressed ETS1 at levels comparable to EWSR1::FLI1-depleted cells (TC-32-ETS1-High) and one expressing lower levels of ETS1 (TC-32-ETS1-Low) (**Fig. 5A, Supplemental Fig. S5A**), neither of which exhibited changes in EWSR1::FLI1 expression **(Fig. 5B)**. Transcriptomic analysis of the TC-32-ETS1-High and ETS1-Low cells versus control cells identified over 5000 genes as exhibiting significantly altered expression in both clones. (**Supplemental Fig. S5C, Supplemental Table S8**). We identified a common set of 1886 downregulated and 1874 upregulated genes in TC-32-ETS1-High and ETS1-Low cells (**Supplemental Fig. S5D; Supplemental Table S8**); however, because the ETS1 protein levels in the TC-32 ETS1-High cells recapitulated most closely those seen upon EWSR1::FLI1 depletion **(Fig. 5A),** we continued our analysis using results generated from this clone, beginning with the identification of genes bound and regulated by ETS1. Intersecting the TC-32 (siFLI1/siNeg) ETS1 ChIP-seq and the ETS1-High mRNA expression datasets, we defined 522 genes as direct targets of ETS1 **(Supplemental Table S8**), including those detailed in **Fig. 5B**, and which included *ATOH8* (**Supplemental Fig. S5E**). Interestingly, of the 265 genes positively regulated by ETS1, 103 also exhibited an increase in expression following depletion of EWSR1::FLI1 in TC-32 cells (**Fig. 5C; Supplemental Table S8**), and of the 257 genes negatively regulated by ETS1, 72 exhibited a decrease in expression (**Supplemental Fig. S5F; Supplemental Table S8**), further suggesting that alterations in ETS1 expression contribute to the EWS gene signature.

**Figure 5:**
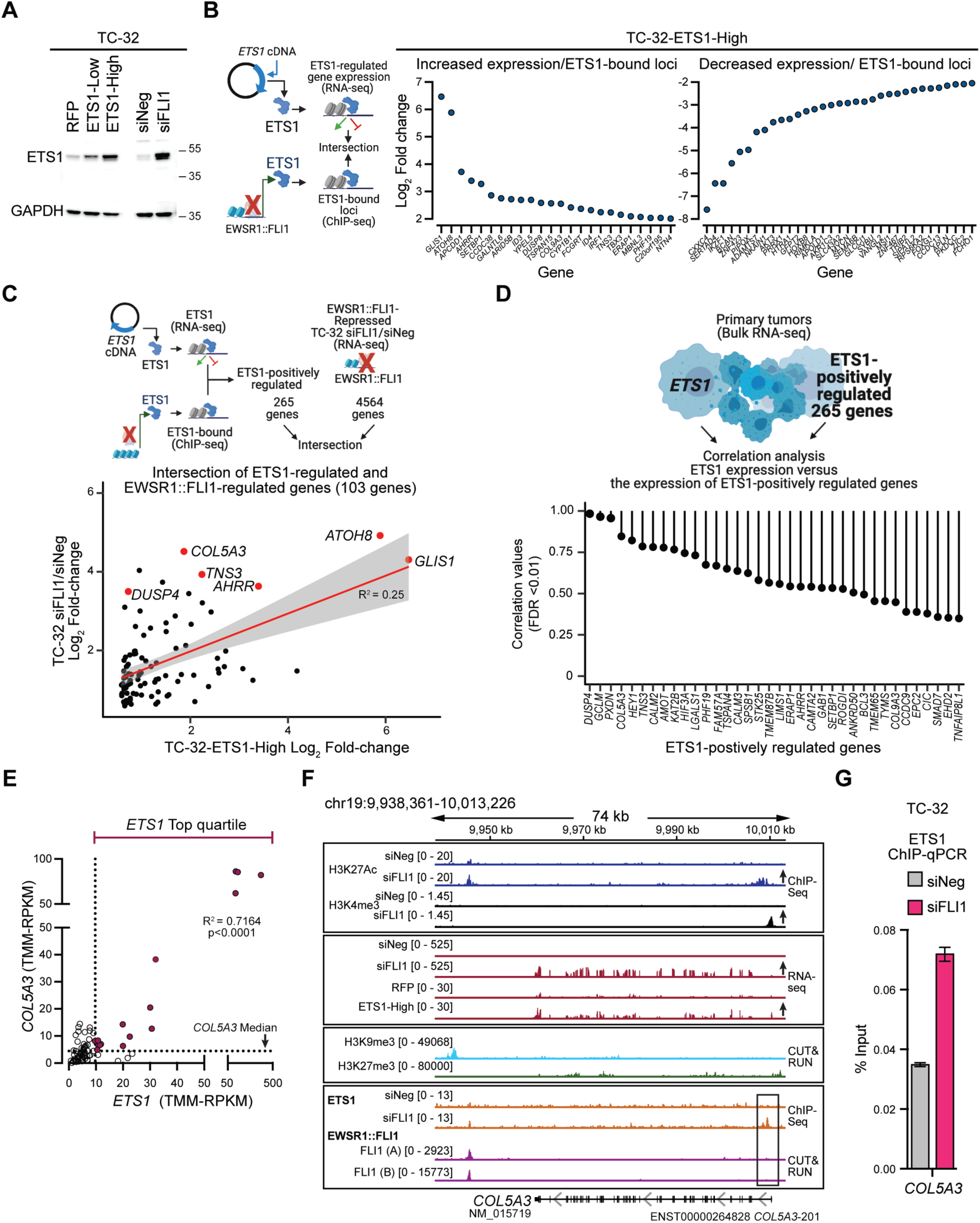
ETS1-regulated genes in EWS cell lines and tumors. **(A)** Immunoblot-based assessment of ETS1 following its ectopic expression (two independent single-cell clones) or upon the depletion of EWSR1::FLI1 in TC-32 cells. **(B)** Model of the intersection of the transcriptome-wide results of ectopic ETS1 expression in TC-32 cells and the detection of ETS1-binding following the depletion of EWSR1::FLI1 in TC-32 cells (left) and results of this analysis focusing on protein-encoding genes that exhibited ±2-fold-change in expression (FDR <0.05) or higher following the ectopic expression of ETS1 (right). **(C)** A model depicting the analysis of the expression of ETS1-positively regulated genes and genes that exhibit an increase in expression following the depletion of the EWSR1::FLI1 and the results of this analysis highlighting selected genes. **(D)** A model depicting the correlation analysis of *ETS1* expression and the expression of ETS1-positively regulated genes in EWS tumors and the results of this analysis using the gene expression profiles of 79 EWS tumors (19) focused on the most significantly correlated genes (FDR <0.01). **(E)** Correlation of the expression of *ETS1* and *COL5A3* in EWS tumor samples (n=79) (19). **(F)** The *COL5A3* locus of TC-32 cells assayed under the indicated conditions showing H3K27ac and H3K4me3 modifications, mRNA, H3K9me3 and H3K27me3 modifications, and ETS1 and EWSR1::FLI1 binding. **(G)** ChIP-qPCR analysis of ETS1 binding at the 5’ end of the *COL5A3* locus in TC-32 cells. Schematics in **B, C,** and **D** were generated using BioRender.

To evaluate the potential contribution of ETS1-regulated genes to EWS, we next assessed their expression levels in EWS tumors relative to *ETS1* expression using the normalized RNA-seq data from 79 EWS tumors (19) by performing a correlation-based statistical analysis of the positively (**Fig. 5D**) and negatively (**Supplemental Fig. S5G**) ETS1-regulated genes. Focusing on the positively regulated ETS1 genes, we observed a correlation between the expression of *ETS1* and 36 of these genes, including *COL5A3*. The positive correlation between *ETS1* and *COL5A3* expression reflects that 16 of the 20 tumors expressing the highest levels of ETS1 (top quartile) also express median or higher levels of *COL5A3* (**Fig. 5E**). Interestingly, three published gene expression datasets (microarray) also show positive correlations of *ETS1* and *COL5A3* expression in EWS tumor samples (R>0.7; **Supplemental Fig. S5H**). ETS1’s putative regulation of *COL5A3* expression is relevant because it encodes one of the type V collagens that forms a component of the ECM (39). Examining the transcription factor binding and epigenetic marks at the *COL5A3* locus and its expression, we observed changes in the H3K27ac and H3K4me3 marks present at the 5’ end of the locus and an increase in its mRNA levels, which the ectopic expression of *ETS1* recapitulated. Under unperturbed conditions, analysis of the annotated repressive marks revealed H3K9me3 deposition in the vicinity of *COL5A3*. Examining ETS1’s putative regulation of *COL5A3* expression, we noted two ETS1 bound sites in the proximity of *COL5A3,* one downstream of its 3’ end that overlaps with an EWSR1::FLI1 binding site and one close to the TSS (**Fig. 5F**), which we confirmed as bound by ETS1 using ChIP-qPCR (**Fig. 5G**). These findings are consistent with our hypothesis that ETS1’s transcriptional activity can promote the expression of genes categorized as repressed by EWSR1::FLI1 and that the activation of one or more these genes have the potential to induce phenotypic changes.

### ETS1-mediated activation promotes EWS cell migration

To investigate the phenotypic consequence of increased ETS1 expression in EWS cells, we activated endogenous *ETS1* expression in SK-N-MC (EWSR1::FLI1) and ES-5838 (EWSR1::ERG) cell lines. We used these cell lines because they originated from tumor cells found at metastatic sites (SK-N-MC—retro-orbital and ES-5838—pleural effusion). However, it is critical to note that after several decades of growth as monolayer cultures, SK-N-MC and ES-5838 cells express comparable levels of their respective fusion oncoproteins as EWS cell lines derived at other stages of disease and sites (**Fig. S6A**) and express little (SK-N-MC) to no (ES-5838) detectable ETS1 (**Supplemental Figs. S6B, S6C**). Following confirmation of ETS1 activation in SK-N-MC and ES-5838 cells (**Supplemental Figs. S6B, S6C**) and determination that expression of their respective EWSR1-fusion oncoproteins remained unaffected (**Supplemental Fig. S6D**), we analyzed their growth and migratory phenotypes. Cell proliferation analysis revealed that endogenous activation of ETS1 did not significantly increase cell growth compared to control cells (unmodified and dCas9-VP64) (**Supplemental Fig. S6E**); however, trans-well chemotaxis assays revealed a significant increase in the migratory phenotype of the ETS1-CRISPR-activated EWS cells compared to control cells (**Fig. 6A**). To further assess the effect of activating ETS1 on the phenotype of SK-N-MC and ES-5838 cells, we next used longitudinal imaging and cell tracking analysis to capture the attachment and movement of cells over 96 hrs. We confirmed minimal differences in the proliferative rate of the control and ETS1-activated cell lines (**Supplemental Fig. S6F**) but observed substantial differences in their movement (**Fig. 6B – 6D**, **Videos 1** – **4**). Specifically, the ETS1-activated cell lines showed a greater range of movement over a defined area than control cells (**Figs. 6C, 6D**), suggesting that ETS1 can regulate EWS cell motility.

**Figure 6:**
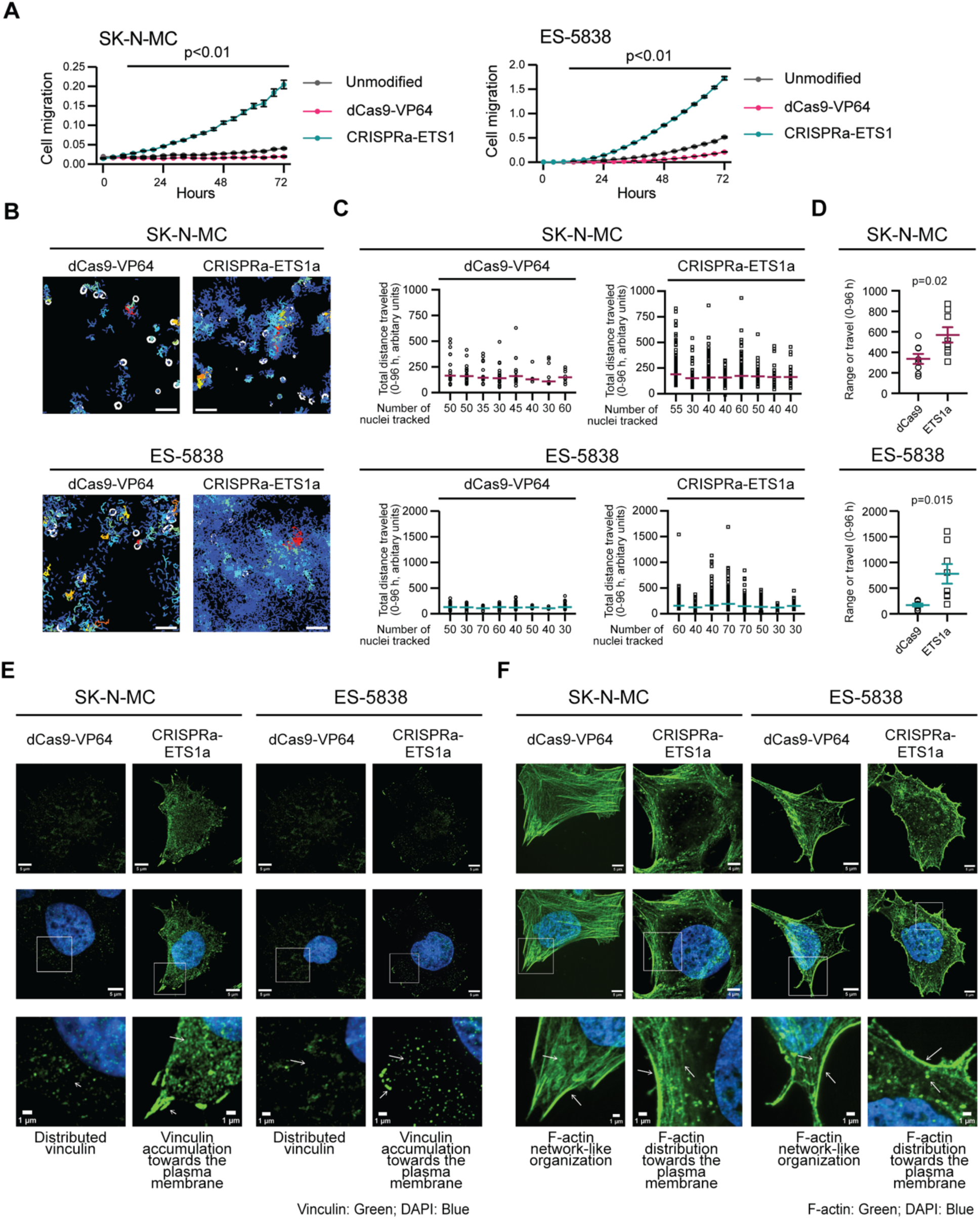
ETS1 promotes the movement of EWS cells. (**A**) Trans-well chemotaxis migration assays with total cell area normalized to the initial top values of the dCas9-VP64 cells in both cell lines (Controls: unmodified and dCas9-VP64 – SK-N-MC 16 replicates each; ES-5838, 32 replicates and CRISPRa-ETS1 EWS cells – SK-N-MC, 24 replicates each, ES-5838, 32 replicates). The results shown represent three independent experiments, and the p-value reports student t-tests performed at each time point. (**B**) Images of the movement of the indicated cells over 96 hrs. See also **Videos 1-4**. (**C**) The quantification of cell movement plotted as total distance traveled over 96 hours (eight ROIs). The first data set in each graph corresponds to the analysis of the ROIs shown in Fig. 6B, and the Y axis indicates the number of nuclei tracked per ROI. (**D**) The range of cell movement was determined for eight ROIs per the indicated cell line shown in Fig. 6C. P-values were determined using an unpaired t-test with Welch’s correction. (**B-D**) Cell movement was determined using images taken every 2 hrs and generated by assessing nuclei (segmented using Otsu thresholds in Fiji) within an ROI of 800 x 800 pixels (one ROI per eight independent wells per cell line) (TrackMate, Fiji plug-in). Assessed nuclei had a mean quality of greater than 100 across all time points with a minimum duration of ten occurrences in 96 hrs as determined by the TrackMate, Fiji plug-in. (**E**) Representative single channel and merged images of vinculin (green) and DAPI (blue) in the indicated cells. Arrows indicate characteristics related to the descriptive statements. (**F**) Representative single channel and merged images of F-actin (green) and DAPI (blue) in the indicated cells. Arrows indicate characteristics related to the descriptive statements included below the zoomed inset. (**E, F**) Cell images, scale bar = 5 µm, zoomed inset, scale bar = 1 µm. Images shown represent 30 images across two independent replicates.

Cell movement involves the formation of focal adhesions and the reorganization of the cytoskeleton. Focal adhesions consist of macromolecular protein complexes that regulate cellular responses to cues from the ECM by controlling mechanical force and downstream signaling cascades that alter the organization of vinculin and F-actin, among other proteins. Using IF and super-resolution microscopy, we compared the distribution of vinculin, (a marker of focal adhesions), and F-actin (a marker of cytoskeletal organization), in the EWS control cells (unmodified and dCas9-VP64) and the cells in which we had activated ETS1 and observed substantial differences (**Figs. 6E, 6F; Supplemental Figs. S6G, S6H**). In control SK-N-MC and ES-5838 cells, we observed low vinculin IF signals distributed throughout cytoplasm (**Fig. 6E, Supplemental Fig. S6G**), but following the activation of ETS1, we observed enhanced vinculin IF signals that accumulated towards the plasma membrane (**Fig. 6D**). Consistent with alterations in the distribution of vinculin, we also observed changes in the organization of F-actin. Specifically, under control conditions, F-actin in SK-N-MC and ES-5838 cells exhibited a network-like organization (**Fig. 6F, Supplemental Fig. S6H**), but a redistribution of F-actin towards the plasma membrane following the activation of ETS1 (**Fig. 6F**). These findings suggest ETS1 regulates the expression of one or more proteins that function in mediating the changes in the configuration of the cytoskeleton and potentially cell movement.

### ETS1 regulates the expression of the focal adhesion protein TENSIN3

Examination of the functions associated with the positively ETS1-regulated genes (**Fig. 5C**), whose expression in tumors also correlated with *ETS1* (**Fig. 5D**), led us to next consider *TNS3* as a possible candidate for contributing to the phenotype of the ETS-activated EWS cells compared with control cells. *TNS3* encodes TENSIN3, a focal adhesion protein recently defined as a regulator of oligodendrocyte differentiation during murine development and the osteogenic differentiation of bone marrow stromal cells (40,41). Not previously studied in EWS, we noted that 18 of 20 EWS tumors (RNA-seq) expressing the highest levels of ETS1 (top quartile) also express median or higher levels of *TNS3* (**Fig. 7A**), and multiple EWS tumor gene expression profiles (microarray-based), show that the expression of *ETS1* and *TNS3* positively correlate (**Fig. 7B, Supplemental Fig. S7A**). Furthermore, high *TNS3* expression is associated with a reduced overall survival probability **(Fig. 7C)**. Under unperturbed conditions, in TC-32 cells, we observed two H3K9me3 peaks in proximity to the 5’ end of *TNS3*, but minimal H3K27me3, H3K4me3, and H3K27ac signals, and no mRNA expression (**Fig. 7D; upper tracks**). Consistent with EWSR1::FLI1’s mediation of this repressive state, we detected EWSR1::FLI1 binding at multiple sites on the *TNS3* gene body (**Fig. 7D; middle purple tracks**). In contrast, following the depletion of EWSR1::FLI1, we observed an increase in the *TNS3* mRNA levels, which *ETS1* ectopic expression, in the presence of EWSR1::FLI1, recapitulated (**Fig. 7D; middle red tracks**). Furthermore, in the absence of EWSR1::FLI1, we observed an increase in the H3K4me3 mark at the TSS of *TNS3,* a concomitant increase in H3K27ac across the locus, and ETS1 binding at a site between exons 1 and 2 of *TNS3* (**Fig. 7D; lower orange tracks**). We also confirmed TNS3’s upregulation in the CRISPRa-ETS1 EWS cell lines using qRT-PCR (**Fig. 7E**), immunoblotting (**Fig. 7F**), and IF (**Fig. 7G**).

**Figure 7:**
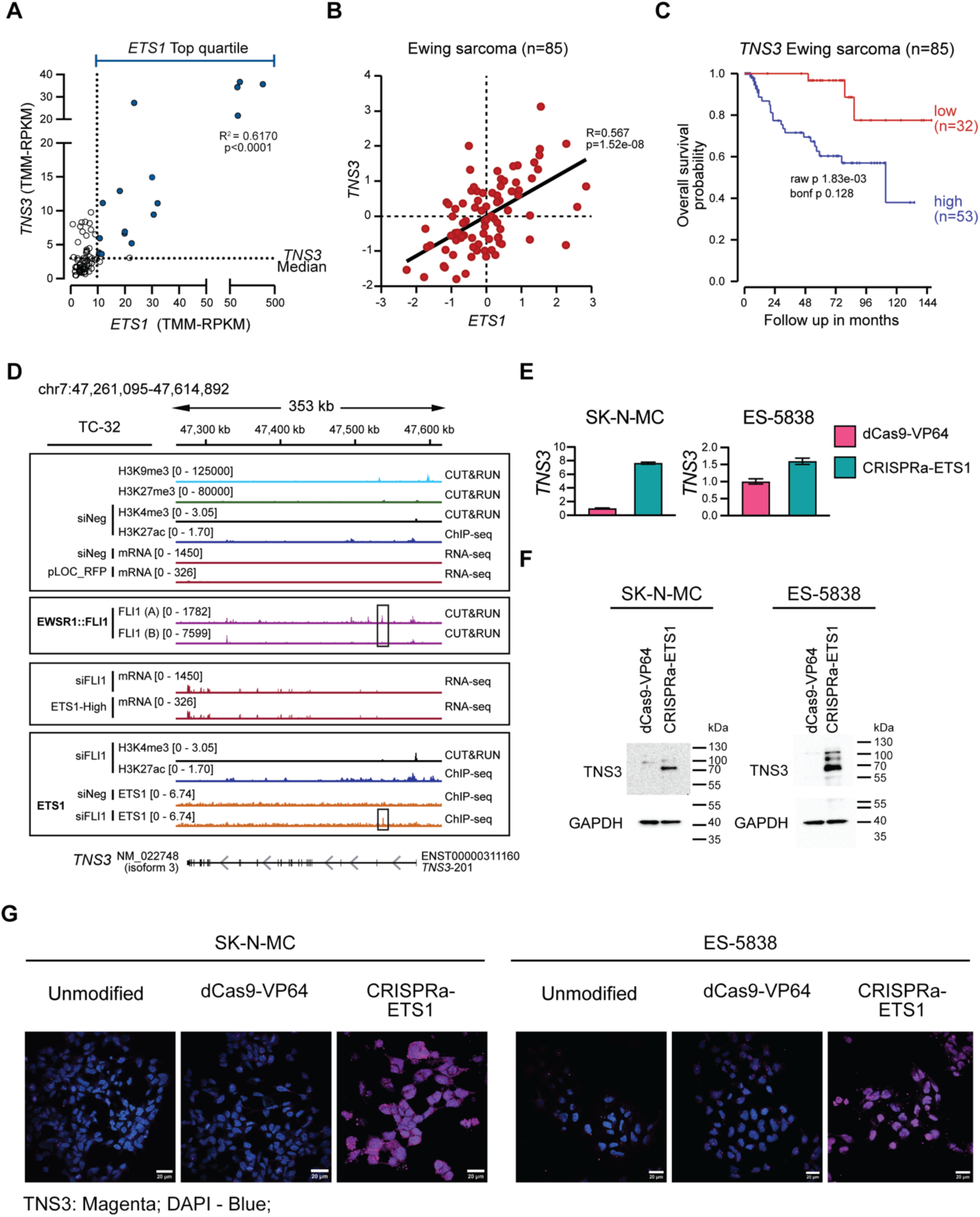
ETS1 regulates the expression of TENSIN3. (**A**) The expression of *ETS1* and *TNS3* in EWS tumor samples (n=79). (**B**) The expression of *ETS1* and *TNS3* mRNA levels (microarray-based analysis) in EWS tumors (n=85) reported previously (21), analyzed and plotted using the R2 Genomics Visualization platform (20). (**C**) Kaplan-Meier curves (overall survival probability) of *TNS3* mRNA levels (microarray-based analysis) in EWS tumors (n=85) reported previously (21) and analyzed and plotted using the R2 Genomics Visualization platform (20). (**D**) The *TNS3* locus of TC-32 cells assayed under control conditions showing H3K9me3, H3K27me3, H3K4me3, and H3K27ac modifications, and mRNA expression and EWSR1::FLI1 binding; H3K4me3 and H3K27ac modifications following the depletion of EWSR1::FLI1; *TNS3* mRNA levels following the depletion of EWSR1::FLI1 or the ectopic expression of *ETS1;* and ETS1 binding in the presence or absence of EWSR1::FLI1. (**E**) The expression of *TNS3* assessed by qRT-PCR analysis in the indicated unmodified and modified EWS cell lines. Results are shown as mean ± SEM of three replicates. (**F**) Immunoblots assessing TNS3 protein expression following the activation of ETS1 in SK-N-MC and ES-5838 cells. **(G)** Confocal images showing the single-channel and merged TNS3 (magenta, IF) and DAPI stain (blue) signals observed in unmodified and modified SK-N-MC or ES-58368 cells as indicated. Scale bar = 25 µm. The images represent 30 images across two independent replicates.

TNS3 contributes to the transduction of signals between the extracellular environment and the cytoskeleton through its binding of focal adhesion complex components, including vinculin, via another cytoskeletal protein, TALIN (42). We thus hypothesized that ETS1’s upregulation of TNS3 contributes to the cytoskeletal reorganization observed in the CRISPRa-ETS1 EWS cells compared to control cells. To evaluate this hypothesis, we used super-resolution microscopy to visualize TNS3 and vinculin in the CRISPRa-ETS1 EWS cells (**Figs 8A, 8B**). Consistent with our previous findings, the control SK-N-MC and ES-5838 cells showed minimal expression of TNS3 and low vinculin IF signals, but following the activation of ETS1, an increase in TNS3 expression and vinculin accumulation at the plasma membrane. Critically, merged images showed evidence of overlapping TNS3 and vinculin signals at specific regions of the plasma membrane, particularly membrane projections. We also examined TNS3’s localization in the context of F-actin organization (**Supplemental Figs. S8A, S8B**). As before, we observed F-actin has a network-like organization in the control cells that becomes distributed towards the plasma membrane upon activation of ETS1. We observed evidence of TNS3 in the proximity of F-actin but minimal overlaps in their signals, suggesting that if TNS3 contributes to cytoskeletal reorganization it is via its adhesion-associated functions (42). To confirm this, we validated two siRNAs targeting *TNS3* (**Supplemental Fig. S8C**) and employed one (siTNS3.2) to assess if depleting TNS3 in the CRISPRa-ETS1 cells alters their phenotype (**Figs. 8C – F**). We observed that a reduction in TNS3 expression in both the CRISPRa-ETS1 cell lines resulted in substantially less regional accumulation of vinculin at the plasma membrane and a reversal of the F-actin phenotype from one of accumulation at the plasma membrane to a more network-like organization. These observations indicate that ETS1-expressing EWS tumor cells have the potential to exhibit TNS3-dependent changes in their cytoskeleton that are consistent with the promotion of cell movement, a critical aspect of the metastatic process.

**Figure 8:**
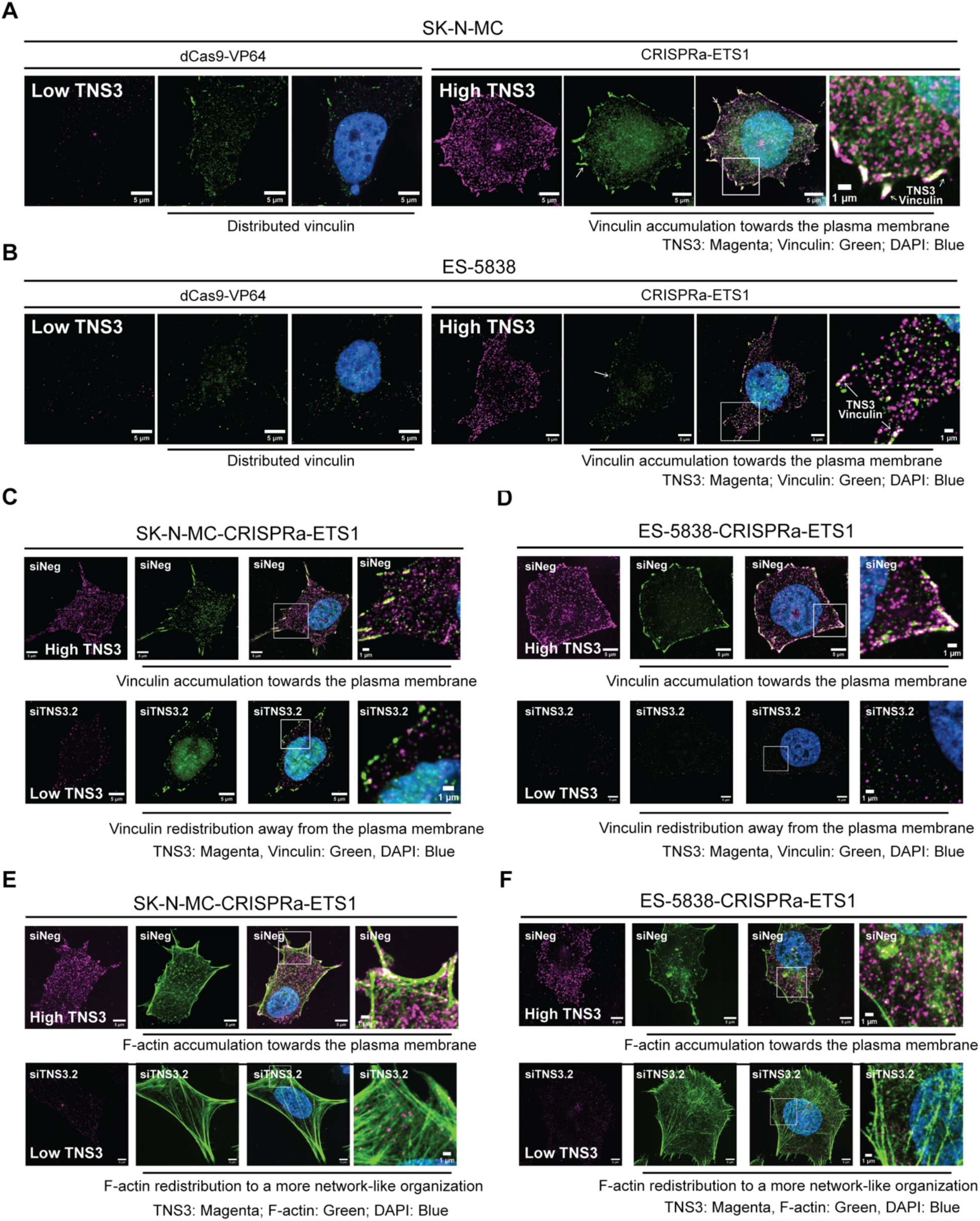
EWS cells expressing ETS1 exhibit TENSIN3-dependent changes in cytoskeletal organization. (**A**) Super-resolution single channel and merged images of TNS3 (magenta, IF), vinculin (green, IF), and DAPI (blue) in the indicated modified SK-N-MC cells, including the indicated inset. (**B**) Super-resolution single channel and merged images of TNS3 (magenta, IF), vinculin (green, IF), and DAPI (blue) in the indicated modified ES-5838 cells, including the indicated inset. (**C, D**) Single channel and merged images (including the indicated inset) of TNS3 (magenta, IF), vinculin (green, IF), and DAPI (blue) in (**C**) the indicated modified SK-N-MC cells and (**D**) the indicated modified ES-5838 cells transfected with control siRNA (siNeg) (upper panels) or a siRNA targeting *TNS3* (lower panels). (**E, F**) Single channel and merged images (including the indicated inset) of TNS3 (magenta, IF), F-actin (green, IF), and DAPI (blue) in (**E**) the indicated modified SK-N-MC cells and (**F**) the indicated modified ES-5838 cells transfected with control siRNA (siNeg) (upper panels) or an siRNA targeting *TNS3* (lower panels). Images represent 30 images across three independent replicates. Cell images, scale bar = 5 µm, insets, scale bar = 1 µm.

## Discussion

Many studies have interrogated the epigenetic mechanisms and genes that the EWSR1::FLI1/ERG fusion proteins activate as these are critical to the initiation and promotion of tumorigenesis (reviewed in 6,43), but less well-characterized is the effect of the altered expression of genes associated with the repressive function of these fusion oncoproteins. However, a convergence of recent reports has demonstrated that the epigenomes of EWS cells *in vivo* are more heterogenous and responsive to external cues than earlier studies had suggested (10,28,44–47). These findings have highlighted the importance of investigating the genes categorized as repressed by EWSR1::FLI1/ERG because of their potential to contribute to processes linked to metastasis if expressed. Here, focusing on transcription factor genes repressed by EWSR1::FLI1 (**Figs. 1** and **2)**, we demonstrated the fusion protein’s repression of the ETS transcription factor, ETS1, *via* the deposition of the polycomb repressive complex-2 (PRC2)-associated H3K27me3 histone modification (**Fig. 3**). Interestingly, though ETS1 and EWSR1::FLI1 both bind the same ETS DNA consensus sequence, we showed that ETS1 and EWSR1::FLI1 occupy distinct regions on chromatin, with the former predominantly binding within promoter regions, and the latter, distal regions (**Fig. 4**). We also demonstrated that ETS1 could function as a transcriptional regulator even in the presence of EWSR1::FLI1, altering the expression of hundreds of genes, including the genes encoding the fibrillar collagen COL5A3 and the focal adhesion protein TNS3, suggesting that ETS1 could exert substantial phenotypic effects if expressed in tumor cells (**Figs. 5** and **7**). Consistent with this concept, we observed that the activation of endogenous ETS1 in two EWS cell lines promotes alterations in their movement (**Fig. 6**) and TNS3-dependent reorganization of the cytoskeleton (**Fig. 8**). Importantly, interrogation of multiple tumor gene expression profiles showed evidence for elevated *ETS1* mRNA level in a subset of Ewing sarcomas, the positive correlation of *ETS1* and *TNS3* expression, and a dataset that enables the correlation gene expression with clinical outcome indicated higher *ETS1* expression is associated with a poorer overall survival probability (**Supplemental Fig. S4A**), as too is the expression of *TNS3*. (**Fig. 7C**).

ETS1 is a critical regulator of normal embryonic development (48,49) that exhibits an entirely different expression pattern in adult tissues when it becomes restricted to immune cell sub-types, including B cells, T cells, and NK cells (50–53). In the developing embryo, ETS1 contributes to the regulation of the mobility and invasive properties of several progenitor cell types, including endothelial cells that contribute to angiogenesis (54–56) and the diverse cell types that arise from neural crest cells (57–63). Interestingly, neural crest cells are one of the few cell-types that tolerate EWSR1::FLI1 expression (64). Studies of mice, chick, and Xenopus embryos have noted *Ets1* expression in pre-migratory and migrating neural crest and neural crest-derived cells, including cardiac and cranial neural crest cells (59–63). The expression of *Ets1* in cardiac neural crest cells appears particularly important as analysis of *Ets1*^-/-^ mice on some genetic backgrounds demonstrate nearly complete perinatal lethality due to the failure of cardiac neural crest cells to migrate appropriately, resulting in heart malformation (60,63). In chick embryos, studies have also defined ETS1’s contributions to the gene regulatory network that defines migratory cranial neural crest cells, which give rise to cartilage and bone, regulating the expression of at least four markers of this cell type, RXRG, LTK, COL9A3, LMO4 (62). Interestingly, we noted ETS1 binding at the *LMO4* and *COL9A3* loci following the silencing of *EWSR1::FLI1* and their upregulation following the ectopic expression of *ETS1* in TC-32 EWS cells (**Supplemental Table S8**). Considering the invasive and migratory characteristics of the progenitor cells that express ETS1, we hypothesized that its expression in an EWS cell could promote the movement of such a cell away from the site of primary tumor growth. Supporting this hypothesis, we observed that the ETS1-CRISPRa EWS cell lines exhibited enhanced cell movement and migration and substantial cytoskeletal reorganization compared to control cells (**Fig. 6**).

The cytoskeletal reorganization that regulates cell movement requires the function of many proteins, including members of the TENSIN protein family (TNS1/2/3), which function as focal adhesion adaptor proteins (reviewed in 65-67). In this study, we noted accumulation of vinculin towards the plasma membrane (**Figs. 6E** and **8A, 8B**), the colocalization of vinculin and TNS3 at membrane projections (**Figs. 8A, 8B** and **8C**, **8D**, upper panels), and the presence of F-actin at the leading edge of cells (**Figs. 6F**, **8E** and **8F**, upper panels, **Supplemental Figs. S8A**, **S8B**); phenotypes consistent with the formation of focal adhesions and cell movement. Critically depletion of TNS3 reversed these changes in the phenotype of ETS1-activated cells to that of cells expressing little or no ETS1 (**Figs. 8E, 8F**, lower panels compared, for example, to the images shown in **Supplemental Fig. S6G** and **S6H**). We present several lines of evidence that ETS1 regulates the expression of TNS3 and that this finding has clinical relevance. First, we observed that ETS1 binds the *TNS3* promoter (**Fig. 7D**) and, second, that the ectopic expression of *ETS1* and activation of endogenous *ETS1* expression increases TNS3 expression (**Figs. 5B, 7E, 7F**). Importantly, interrogation of tumor gene expression profiles from five independent datasets showed a positive correlation of *ETS1* and *TNS3* expression (**Figs. 7A** and **7B**, and **Supplemental Fig. S7A**) and association of high *TNS3* expression with a poorer overall survival probability (**Fig. 7C**). Building on these observations, critical next steps will include further assessment of the phenotypic consequences of TNS3 expression in EWS cells and determination of whether elevated expression of ETS1, TNS3, or both contribute to the risk of metastasis.

Future studies will also need to investigate if the expression of ETS1 and the other transcription factor genes data suggests some EWS tumor cells express at higher levels than EWS cell lines is a result of heterogenous EWSR1:FLI1/ERG expression and/or due to the transduction of external stimuli. The mechanisms by which EWSR1::FLI1 represses gene expression in cell lines and to what extent this is recapitulated in tumors are less well-defined than its recruitment of factors that activate gene expression, but previous studies have shown that in contrast to gene activation, the repression of genes by EWSR1::FLI1 occurs at non-repeat canonical ETS binding sites (7). Here, we identified H3K27Me3 and/or H3K9Me3 marks adjacent to a substantial number of the transcription factor genes repressed by EWSR1::FLI1/ERG (**Fig. 2**, **3**), though not all, suggesting that EWSR1-fusion proteins can mediate gene repression through additional mechanisms. H3K27me3 repression is facultative and allows repressed genes to be accessible by transcription factor binding in a different cellular state, including during development (reviewed in 68,69). Interestingly, we observed H3K27me3 marks adjacent to *ETS1* and *SNAI2,* both of which exhibited increased expression following depletion of EWSR1::FLI1 (**Fig. 3B, 3D**, **Fig 2G**), confirming that at these loci, fusion oncoprotein activity does not mediate the formation of an irreversible heterochromatic state. It is noteworthy that a recent comprehensive study of 18 EWS cell lines that examined their expression profiles using high-density microarrays under control conditions and following the shRNA-mediated targeting of *EWSR1::FLI1* or *EWSR1::ERG* also reported the upregulation of *SNAI2* in 16 of 18 cell lines (70). However, a further comparison of our observations and the datasets reported by Orth and coworkers detected only slight increases in *ETS1* expression in A673 and TC-32 cells and no change in *ETS1* expression in TC-71, SK-N-MC, and TC-106 cells, among others. It is unclear whether these differences reflect the relative sensitivity of the different assays used. Nevertheless, we have reproducibly detected changes in *ETS1* expression by RNA-seq and/or qPCR and protein analysis following siRNA-mediated targeting *of EWSR1-FLI1*/*ERG* fusion transcripts in multiple EWS cell lines (**Fig. 1**) and found evidence that at least some EWS tumors express ETS1. Critically, we show that even in the presence of EWSR1::FLI1, the expression of ETS1 could have profound effects on the phenotype of EWS cells, including the promotion of cell movement, an essential aspect of the metastatic process.

## Acknowledgments

The Intramural Research Program of the National Cancer Institute (NCI), Center for Cancer Research (CCR), National Institutes of Health supported this study: ZIA BC 011704 supports N.J.C; ZIC BC 010858 supports the CCR Confocal Microscopy Core Facility; and the generation of CRISPR-Cas9 reagents included the support of Federal funds from the National Cancer Institute, National Institutes of Health, under Contract No. HHSN261201500003I. This project was funded in whole or in part with Federal funds from the National Cancer Institute, National Institutes of Health, Department of Health and Human Services, under Contract No. 75N91019D00024. This work utilized the computational resources of the NIH HPC Biowulf cluster (http://hpc.nih.gov). We thank Raj Chari and the Genome Modification Core, Laboratory Animal Sciences Program at the Frederick National Lab for Cancer Research for assistance with the generation of CRISPR-Cas9 reagents, and the CCR Flow Cytometry and CCR Sequencing Cores for technical assistance, and the CCR Microscopy Core, particularly Michael Kruhlak and Andy Tran for technical support and advice. We thank Javed Khan, Jun S. Wei, and Young Song for access to Ewing sarcoma RNA-seq data (Oncogenomics Section, Genetics Branch). We thank Carla Neckles and Alison Cross, former members of the Functional Genetics Section, Genetics Branch, for early discussion and technical assistance related to this work, and Sanjit Mukherjee, Bob Walker, and Fan Yang (Molecular Genetics Section, Genetics Branch) for advice and guidance. We also thank Christine Heske, Pediatric Oncology Branch, CCR for useful discussion.

The content of this publication does not necessarily reflect the views or policies of the U.S. Department of Health and Human Services, nor does mention of trade names, commercial products, or organizations imply endorsement by the U.S. Government.

## Videos, Supplemental Tables and Figures

### Videos

**Video 1:** SK-N-MC-dCas9-VP64

**Video 2:** SK-N-MC-CRISPRa-ETS1

**Video 3:** ES-5838-dCas9-VP64

**Video 4:** ES-5838-CRISPRa-ETS1

### Supplemental Tables

**Supplemental Table S1:** Reagents and resources

**Supplemental Table S2:** RNA-seq analysis of EWSR1::FLI1-depleted EWS cells

**A:** TC-32 cells - RNA-seq analysis following depletion of EWSR1::FLI1

**B:** TC-71 cells - RNA-seq analysis following depletion of EWSR1::FLI1

**C:** A673 cells - RNA-seq analysis following depletion of EWSR1::FLI1

**D:** Increased expression following depletion of EWSR1::FLI1; TC-32, TC-71, and A673

**E:** Decreased expression following depletion of EWSR1::FLI1; TC-32, TC-71, and A673

**F:** Gene ontology analysis - Increased expression following depletion of EWSR1::FLI1

**G:** Gene ontology analysis - Decreased expression following depletion of EWSR1::FLI1

**Supplemental Table S3:** EWSR1::FLI1-regulated transcription factor genes

**A:** Transcription factor genes - Increased expression following depletion of EWSR1::FLI1 (Repressed EWSR1::FLI1 transcription factor (TF) genes)

**B:** Transcription factor genes - Decreased expression following depletion of EWSR1::FLI1 (Activated EWSR1::FLI1 transcription factor (TF) genes)

**Supplemental Table S4:** The epigenomic analysis of TC-32 EWS cells

**A**: TC-32 FLI1 CUT&RUN A

**B:** TC-32 FLI1 CUT&RUN B

**C:** TC-32 H3K9me3 CUT&RUN

**D:** TC-32 H3K27me3 CUT&RUN

**E:** TC-32 H3K27ac CUT&RUN

**F:** TC-32 H3K27ac siNeg ChIP

**G:** TC-32 H3K27ac siFLI1 ChIP

**Supplemental Table S5:** The epigenomic analysis of transcription factor genes repressed by

EWSR1::FLI1

**A:** Transcription factor genes: Intersection of EWSR1::FLI1 binding, H3K9me2, H3K27Me3

**B**: Transcription factor genes: TC-32 FLI1 CUT&RUN A

**C:** Transcription factor genes: TC-32 FLI1 CUT&RUN B

**D:** Transcription factor genes: TC-32 H3K9me3 CUT&RUN

**E:** Transcription factor genes: TC-32 H3K27me3 CUT&RUN

**F:** Transcription factor genes: EWS overall survival probability parameters extracted from the Ewing sarcoma tumor gene expression dataset detailed in (21) using the R2 Genomics Visualization platform (20)

**Supplemental Table S6:** Analysis of H3K4me3 modifications

**A:** TC-32 H3K4me3 siNeg ChIP

**B:** TC-32 H3K4me3 siFLI1 ChIP

**Supplemental Table S7:** Analysis of ETS1 binding

TC-32 ETS1 ChIP-seq siFLI1/siNeg

**Supplemental Table S8:** Analysis of ETS1 ectopic expression

**A:** Differential gene expression analysis of TC-32-pLOC-ETS1-Low versus TC-32-pLOC-RFP

**B:** Differential gene expression analysis of TC-32-pLOC-ETS1-High versus TC-32 pLOC-RFP

**C:** The intersection of TC-32-pLOC-ETS1-Low and TC-32-pLOC-ETS1-High differentially expressed genes

**D:** Genes exhibiting increased expression following the ectopic expression of ETS1 and bound by ETS1 (TC-32 cells)

**E:** Genes exhibiting decreased expression following the ectopic expression of ETS1 and bound by ETS1 (TC-32 cells)

**F:** Genes exhibiting increased expression following the ectopic expression of ETS1, bound by ETS1, and that exhibit an increase in expression following the depletion of EWSR1::FLI1 (TC-32 cells)

**G:** Genes exhibiting decreased expression following the ectopic expression of ETS1, bound by ETS1, and that exhibit a decrease in expression following the depletion of EWSR1::FLI1 (TC-32 cells)

### Supplemental Figures

**Figure S1: The expression profiles of EWSR1::FLI1-depleted cells (A)** qRT-PCR-based validation of the silencing of *EWSR1::FLI1* in the indicated cell lines (mean ± SEM from three independent experiments; p-values determined using unpaired t-test). **(B)** Changes in gene expression were observed following the silencing of *EWSR1::FLI1* in the indicated cell lines. **(C)** Gene ontology analysis of the significantly up-regulated (1107) or down-regulated (1111) genes observed following the silencing of *EWSR1::FLI1* in three EWS cell lines (TC-32, TC-71, and A673). Results are shown for those GO terms that appeared in both the Metascape and GSEA databases (https://metascape.org/gp/index.html-/main/step1 and https://www.gsea-msigdb.org/gsea/msigdb/) plotting the P values from the Metascape analysis. **(D)** RNA-seq analysis of transcription factors genes that demonstrated a significant decrease in expression following the silencing of *EWSR1::FLI1* in TC-32, TC-71, or A673 (three biological replicates; fold change <-1.5, FDR < 0.05).

**Figure S2: EWSR1::FLI1 repressed gene targets defined using EWS cell lines exhibit a broader range of expression in EWS primary tumors. (A)** Comparative expression of selected transcription factor genes in EWS tumor samples (n=79) and EWS cell lines (n=42) (19) (p-values determined using an unpaired t-test with Welch’s correction). **(B)** qRT-PCR-based validation of the silencing of *EWSR1::FLI1* in SK-N-MC and *EWSR1::ERG* in TC-106 cells (mean ± SEM from three independent experiments; p-values determined using unpaired t-test).

**Figure S3: The genome-wide binding of EWSR1::FLI1. (A)** Genome-wide heatmaps of H3K27ac ChIP-seq peak-centered signals in control (siNeg) and *EWSR1::FLI1*-silenced (siFLI1) TC-32 cells showing 10kb windows. **(B)** DNA motif enrichment analysis of EWSR1::FLI1 binding sites determined by CUT&RUN sequence analysis (two independent datasets, A and B, two biological replicates each). (**C)** Clustered heatmaps of ChIP-seq peak-centered signals for EWSR1::FLI1 and H3K27ac in TC-32 cells showing 5kb windows. Regions ranked by EWSR1::FLI1 signals with gene promoter regions defined as – 3kb and + 1kb from TSS. Distal regions were defined as existing outside of promoter regions. **(D)** The *CCND1* locus of TC-32 cells showing the EWSR1::FLI1 and H3K27ac peaks, H3K27ac ChIP-seq signals (TC-32 siNeg or siFLI1-transfected), and H3K9me3 and H3K27me3 signals. **(E)** The *PHLDA1* locus of TC-32 cells showing the EWSR1::FLI1 and H3K27ac peaks, H3K27ac ChIP-seq signals (TC-32 siNeg or siFLI1-transfected), and H3K9me3 and H3K27me3 signals.

**Figure S4: ETS1 is a repressed target of EWSR1::FLI1 in EWS cells.** (**A**) The overall survival probability data associated with *ETS1* expression was extracted from GSE63157 (85 tumors) (21) using the R2 Genomics Analysis and Visualization Platform (20) and plotted as Kaplan Meier curves based on the log-rank. (**B**) EWSR1::FLI1 binding sites at the *ETS1* locus and the sequences at the indicating regions used as the basis of qPCR assays reported in Fig. 3. The underlined sequences indicate the amplified regions and the bolded GGAA and TTCC sequences indicate the binding motifs associated with ETS transcription factors. (**C**) H3K27me deposition at the *ETS1* locus and the sequences at the indicating regions used as the basis of qPCR assays reported in Fig. 3. (**D**) Genome-wide heatmaps of H3K4me3 ChIP-seq peak-centered signals in control (siNeg) and *EWSR1::FLI1*-silenced (siFLI1) TC-32 cells showing 10kb windows.

**Figure S5: Ectopic expression of ETS1 has transcriptome-wide effects.** (**A**) qRT-PCR analysis of *ETS1* expression exhibited by the control and the two single-cell clones expressing an *ETS1* cDNA (NM_01143820; isoform 1). Results are shown as mean ± SEM of three replicates. (**B**) Immunoblots of whole-cell lysates prepared from the indicated cell lines were analyzed using antibodies against the indicated proteins. (**C**) Changes in gene expression were observed following the ectopic expression of *ETS1*. (**D**) The co-occurrence of genes that exhibit significant changes in expression following the ectopic expression of *ETS1* in the indicated TC-32 single-cell clones (see also **Supplemental Table S8**). (**E**) The *ATOH8* locus of TC-32 cells showing ETS1 ChIP-seq peaks (siNeg or siFLI1-transfected) and RNA-seq read depth (siNeg or siFLI1-transfected cells or modified – RFP or ETS1-High). **(F)** A model depicting the analysis of the expression of ETS1-negatively regulated genes and genes that exhibit a decrease in expression following the depletion of EWSR1::FLI1 and the result of this analysis is presented as a scatter plot. **(G)** A model depicting the correlation analysis of *ETS1* expression and the expression of ETS1-negatively regulated genes in EWS tumors and the results of this analysis using the gene expression profiles of 79 EWS tumors (19) focused on significantly correlated genes (FDR <0.05). (**H**) The correlation of *ETS1* and *COL5A3* mRNA levels in EWS tumors reported previously in the indicated studies: GSE142162 79 samples (24); GSE34620 (117 tumors) (22); GSE12102 (37 tumors) (23), analyzed and plotted using the R2 Genomics Visualization platform (20).

**Figure S6: ETS1 promotes the migration of EWS cells.** (**A**) Immunoblots of whole-cell lysates prepared from the indicated cell lines and analyzed using antibodies against the indicated proteins. (**B**) The expression of *ETS1* assessed by qRT-PCR analysis in the indicated unmodified and modified EWS cell lines. Results are shown as mean ± SEM of three replicates, except for duplicate ES-5838-dCas9-VP64 samples. (**C**) Immunoblot analysis of whole-cell lysates prepared from the indicated unmodified and modified EWS cell lines and analyzed using the antibodies against the indicated proteins. (**D**) Immunoblot analysis of whole-cell lysates prepared from the indicated unmodified and modified EWS cell lines and analyzed using the antibodies against the indicated proteins. (**E**) Cell confluency (area of confluency normalized to 0 hrs) over time of the indicated unmodified and modified (dCas9-VP64 and CRISPRa-ETS1) EWS cell lines. Each data point indicates the mean ± SEM of four replicates representing three independent experiments. (**F**) Cell confluency (area of confluency normalized to 0 hrs) over time of dCas9-VP64 and CRISPRa-ETS1 EWS cell lines employed in the analysis of cell movement (**Fig. 6B, 6C** and **videos 1** - **4**). The data shown indicates the mean ± SEM of 46 replicates. (**G)** Super-resolution merged images of unmodified SK-N-MC and ES-5838 cells analyzed using an antibody against vinculin (green) and the DAPI stain (blue). (**H**) Super-resolution merged images of unmodified SK-N-MC and ES-5838 cells analyzed using an antibody against F-actin (green) and the DAPI stain (blue).

**Figure S7: ETS1 regulates the expression of TENSIN3.** (**A**) The correlation of *ETS1* and *TNS3* mRNA levels in EWS tumors reported previously in the indicated studies: GSE142162 79 samples (24); GSE34620 (117 tumors) (22); GSE12102 (37 tumors) (23), analyzed and plotted using the R2 Genomics Visualization platform (20). (**B**) Confocal single-channel images of DAPI (blue) and TNS3 (magenta) in unmodified and modified SK-N-MC cells corresponding to the merged images shown in Fig. 7G. (**C**) Confocal single-channel images of DAPI (blue) and TNS3 (magenta) in unmodified and modified ES-5838 cells corresponding to the merged images shown in Fig. 7G.

**Figure S8: EWS cells expressing ETS1 exhibit TENSIN3-dependent changes in cytoskeletal organization.** (**A**) Super-resolution single-channel and merged images (including a zoomed inset) of the indicated modified SK-N-MC cells analyzed using antibodies against TNS3 (magenta), F-Actin (green), and the DAPI stain (blue). (**B**) Super-resolution single-channel and merged images (including a zoomed inset) of the indicated modified ES-5838 cells analyzed using antibodies against TNS3 (magenta), F-Actin (green), and the DAPI stain (blue). (**A, B**) Cell images, scale bar = 5 µm, zoomed inset, scale bar = 1 µm; images represent 30 images across two independent replicates. (**C**) Immunoblots of whole-cell lysates prepared from cells transfected with the indicated siRNAs for 48 hrs and analyzed using the antibodies against the indicated proteins. Studies examining TNS3 function in mediating changes in the cytoskeletal organization used the siRNA designated siTNS3.2 (**Fig. 8C-F**).

## Authors’ Contributions

**Conception and design:** Vernon Justice Ebegboni and Natasha J. Caplen

**Development of methodology:** Vernon Justice Ebegboni, Soumya Sundara Rajan, Tamara L. Jones

**Acquisition of data:** Vernon Justice Ebegboni, Soumya Sundara Rajan, Tamara L. Jones, Tayvia Brownmiller

**Analysis and data interpretation (e.g., statistical analysis, biostatistics, computational analysis):** Vernon Justice Ebegboni, Erica C. Pehrsson, Patrick X. Zhao, Natasha J. Caplen

**Writing, review, and/or revision of the manuscript:** Vernon Justice Ebegboni, Soumya Sundara Rajan, Natasha J. Caplen, Tamara L. Jones, Tayvia Brownmiller, Erica C. Pehrsson

**Administrative, technical, or material support (i.e., reporting or organizing data, constructing databases):** Tamara L. Jones, Vernon Justice Ebegboni, Erica C. Pehrsson, Patrick X. Zhao

**Study supervision:** Natasha J. Caplen

