## Supplemental Figures, Methods, Table S1 for "ETS1, a target gene of the EWSR1::FLI1 fusion oncoprotein, regulates the expression of the focal adhesion protein TENSIN3"

**Supplemental Materials and Methods**

**Supplemental Table 1**

**Figure S1**

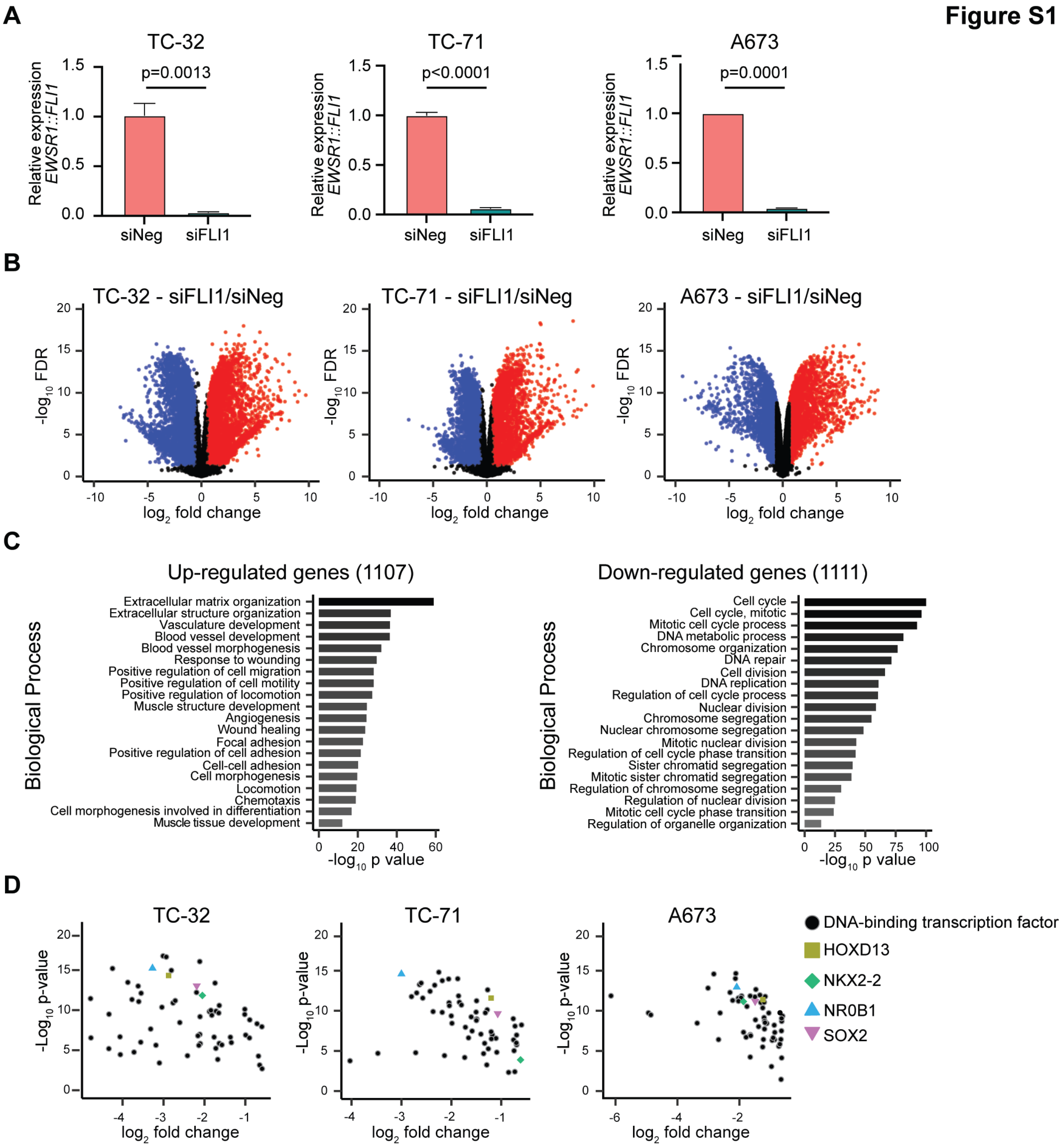

**Figure S1: The expression profiles of EWSR1::FLI1-depleted cells (A)** qRT-PCR-based validation of the silencing of *EWSR1::FLI1* in the indicated cell lines (mean ± SEM from three independent experiments; p-values determined using unpaired t-test). **(B)** Changes in gene expression were observed following the silencing of *EWSR1::FLI1* in the indicated cell lines. **(C)** Gene ontology analysis of the significantly up-regulated (1107) or down-regulated (1111) genes observed following the silencing of *EWSR1::FLI1* in three EWS cell lines (TC-32, TC-71, and A673). Results are shown for those GO terms that appeared in both the Metascape and GSEA databases (https://metascape.org/gp/index.html - /main/step1 and https://www.gsea-msigdb.org/gsea/msigdb/) plotting the P values from the Metascape analysis. **(D)** RNA-seq analysis of transcription factors genes that demonstrated a significant decrease in expression following the silencing of *EWSR1::FLI1* in TC-32, TC-71, or A673 (three biological replicates; fold change <-1.5, FDR < 0.05).

**Figure S2**

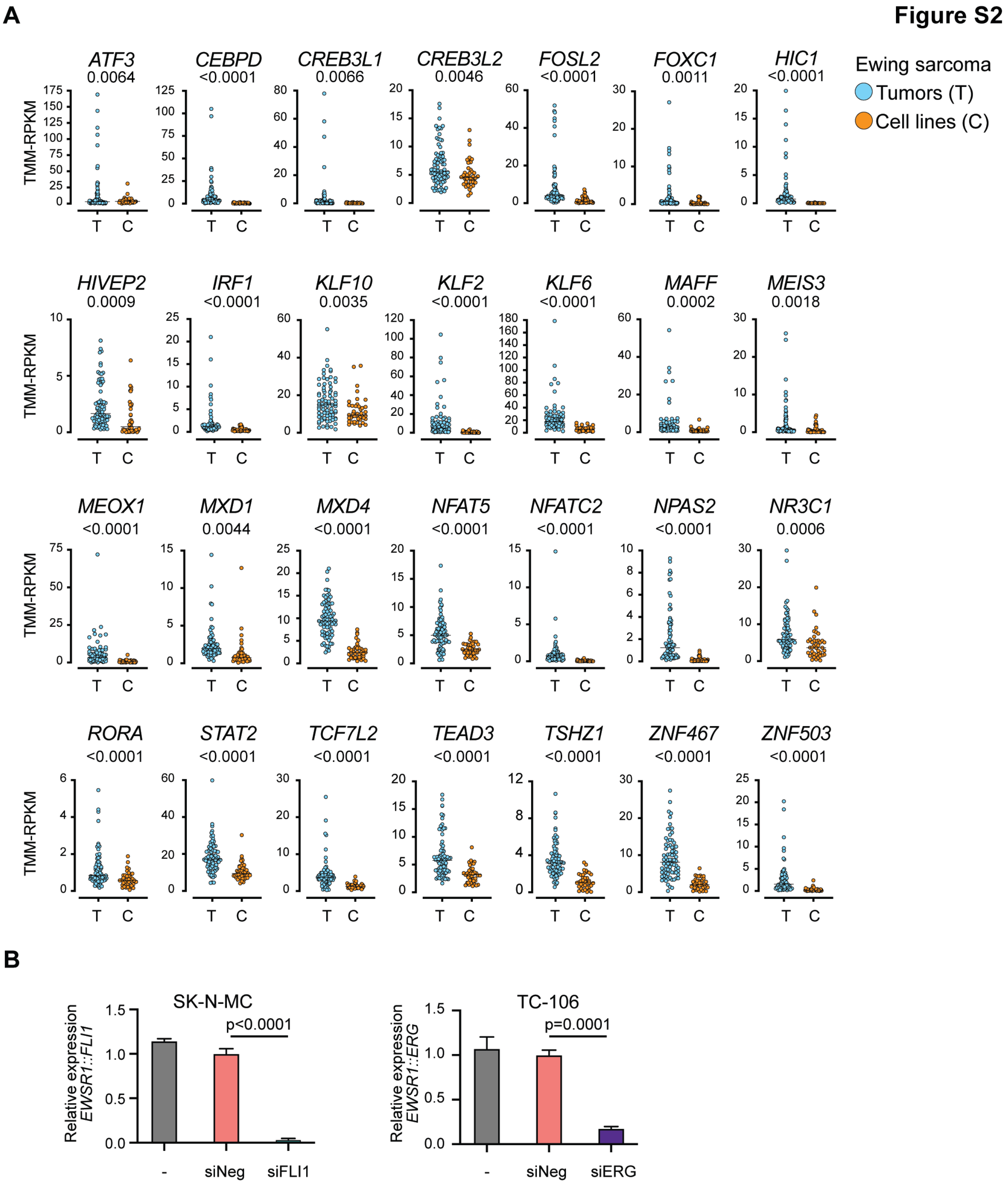

**Figure S2: EWSR1::FLI1 repressed gene targets defined using EWS cell lines exhibit a broader range of expression in EWS primary tumors. (A)** Comparative expression of selected transcription factor genes in EWS tumor samples (n=79) and EWS cell lines (n=42) (19) (p-values determined using an unpaired t-test with Welch's correction). **(B)** qRT-PCR-based validation of the silencing of *EWSR1::FLI1* in SK-N-MC and *EWSR1::ERG* in TC-106 cells (mean ± SEM from three independent experiments; p-values determined using unpaired t-test).

**Figure S3**

A

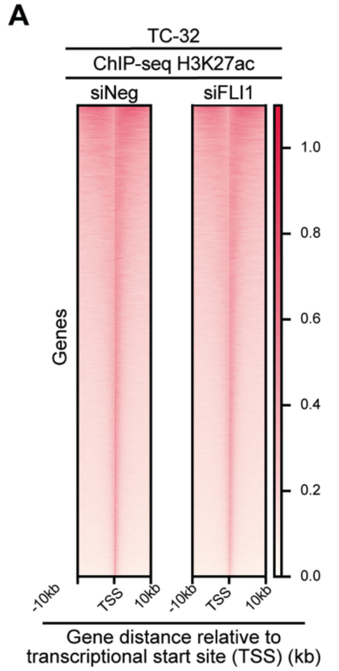

B

TC-32 CUT&RUN (A) 40617 EWSR1::FLI1 peaks

| | Transcription factor | Motif | p value $-\log_{10}$ |
|---|---|---|---|
| 1 | FLI1 (ETS) | | 6.653e+03 |
| 2 | ELK1 (ETS) | | 6.153e+03 |
| 3 | EWS:FLI1 (ETS) | | 6.106e+03 |
| 4 | ELK4 (ETS) | | 5.991e+03 |
| 5 | ETV4 (ETS) | | 5.910e+03 |
| 6 | GABPA1 (ETS) | | 5.887e+03 |
| 7 | ETV1 (ETS) | | 5.177e+03 |
| 8 | ETV2 (ETS) | | 5.115e+03 |
| 9 | ETS1 (ETS) | | 4.612e+03 |
| 10 | ERG (ETS) | | 4.265e+03 |

TC-32 CUT&RUN (B) 38221 EWSR1::FLI1 peaks

| | Transcription factor | Motif | p value $-\log_{10}$ |
|---|---|---|---|
| 1 | FLI1 (ETS) | | 5.285e+03 |
| 2 | ETV4 (ETS) | | 4.644e+03 |
| 3 | ETV2 (ETS) | | 4.443e+03 |
| 4 | GABPA (ETS) | | 4.369e+03 |
| 5 | ETV1 (ETS) | | 4.265e+03 |
| 6 | EWS:FLI1 (ETS) | | 4.064e+03 |
| 7 | ERG (ETS) | | 3.807e+03 |
| 8 | ETS1 (ETS) | | 3.738e+03 |
| 9 | ELK1 (ETS) | | 3.714e+03 |
| 10 | EWS:ERG (ETS) | | 3.402e+03 |

C

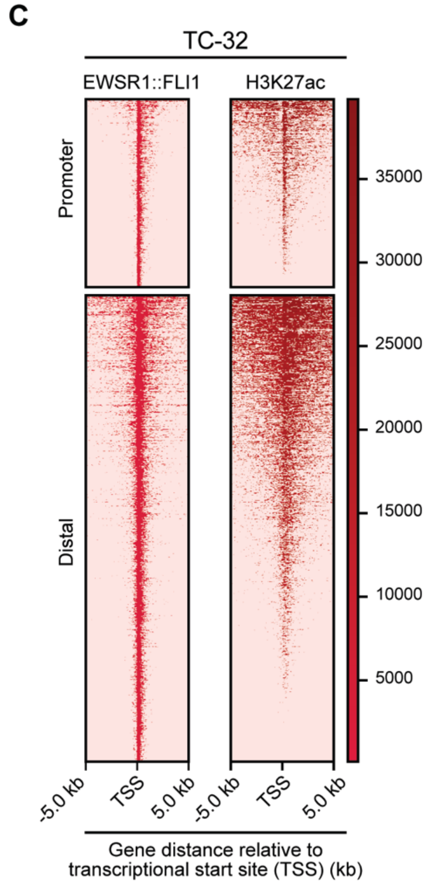

D

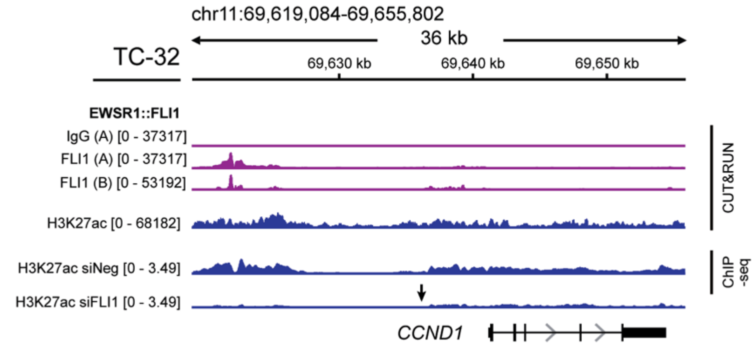

E

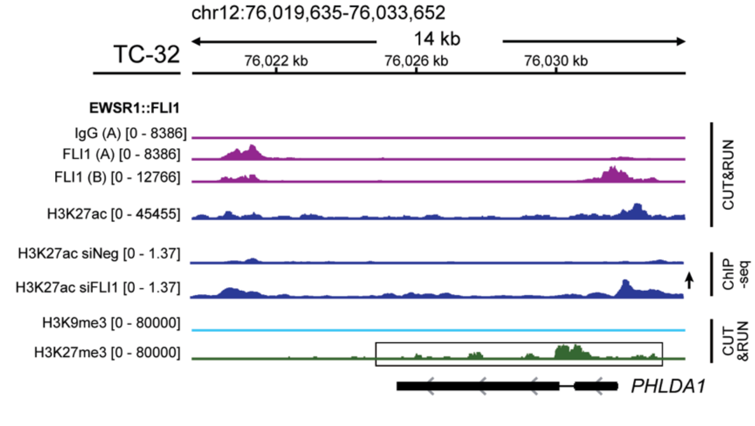

**Figure S3: The genome-wide binding of EWSR1::FLI1. (A)** Genome-wide heatmaps of H3K27ac ChIP-seq peak-centered signals in control (siNeg) and *EWSR1::FLI1*-silenced (siFLI1) TC-32 cells showing 10kb windows. **(B)** DNA motif enrichment analysis of EWSR1::FLI1 binding sites determined by CUT&RUN sequence analysis (two independent datasets, A and B, two biological replicates each). (**C**) Clustered heatmaps of ChIP-seq peak-centered signals for EWSR1::FLI1 and H3K27ac in TC-32 cells showing 5kb windows. Regions ranked by EWSR1::FLI1 signals with gene promoter regions defined as – 3kb and + 1kb from TSS. Distal regions were defined as existing outside of promoter regions. **(D)** The *CCND1* locus of TC-32 cells showing the EWSR1::FLI1 and H3K27ac peaks, H3K27ac ChIP-seq signals (TC-32 siNeg or siFLI1-transfected), and H3K9me3 and H3K27me3 signals. **(E)** The *PHLDA1* locus of TC-32 cells showing the EWSR1::FLI1 and H3K27ac peaks, H3K27ac ChIP-seq signals (TC-32 siNeg or siFLI1-transfected), and H3K9me3 and H3K27me3 signals.

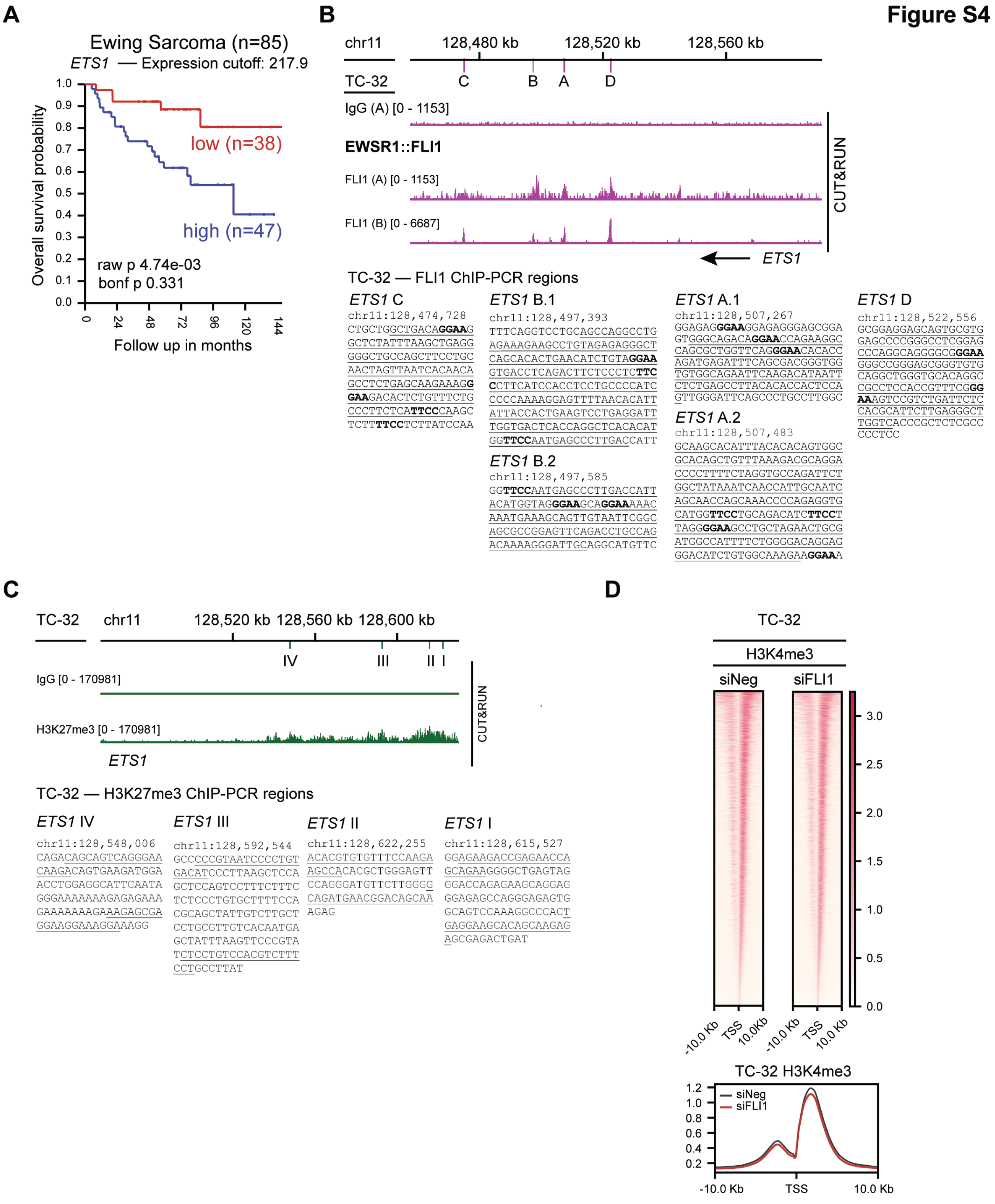

**Figure S4: ETS1 is a repressed target of EWSR1::FLI1 in EWS cells.** (**A**) The overall survival probability data associated with *ETS1* expression was extracted from GSE63157 (85 tumors) (21) using the R2 Genomics Analysis and Visualization Platform (20) and plotted as Kaplan Meier curves based on the log-rank. (**B**) EWSR1::FLI1 binding sites at the *ETS1* locus and the sequences at the indicating regions used as the basis of qPCR assays reported in **Fig. 3**. The underlined sequences indicate the amplified regions and the bolded GGAA and TTCC sequences indicate the binding motifs associated with ETS transcription factors. (**C**) H3K27me deposition at the *ETS1* locus and the sequences at the indicating regions used as the basis of qPCR assays reported in **Fig. 3**. (**D**) Genome-wide heatmaps of H3K4me3 ChIP-seq peak-centered signals in control (siNeg) and *EWSR1::FLI1*-silenced (siFLI1) TC-32 cells showing 10kb windows.

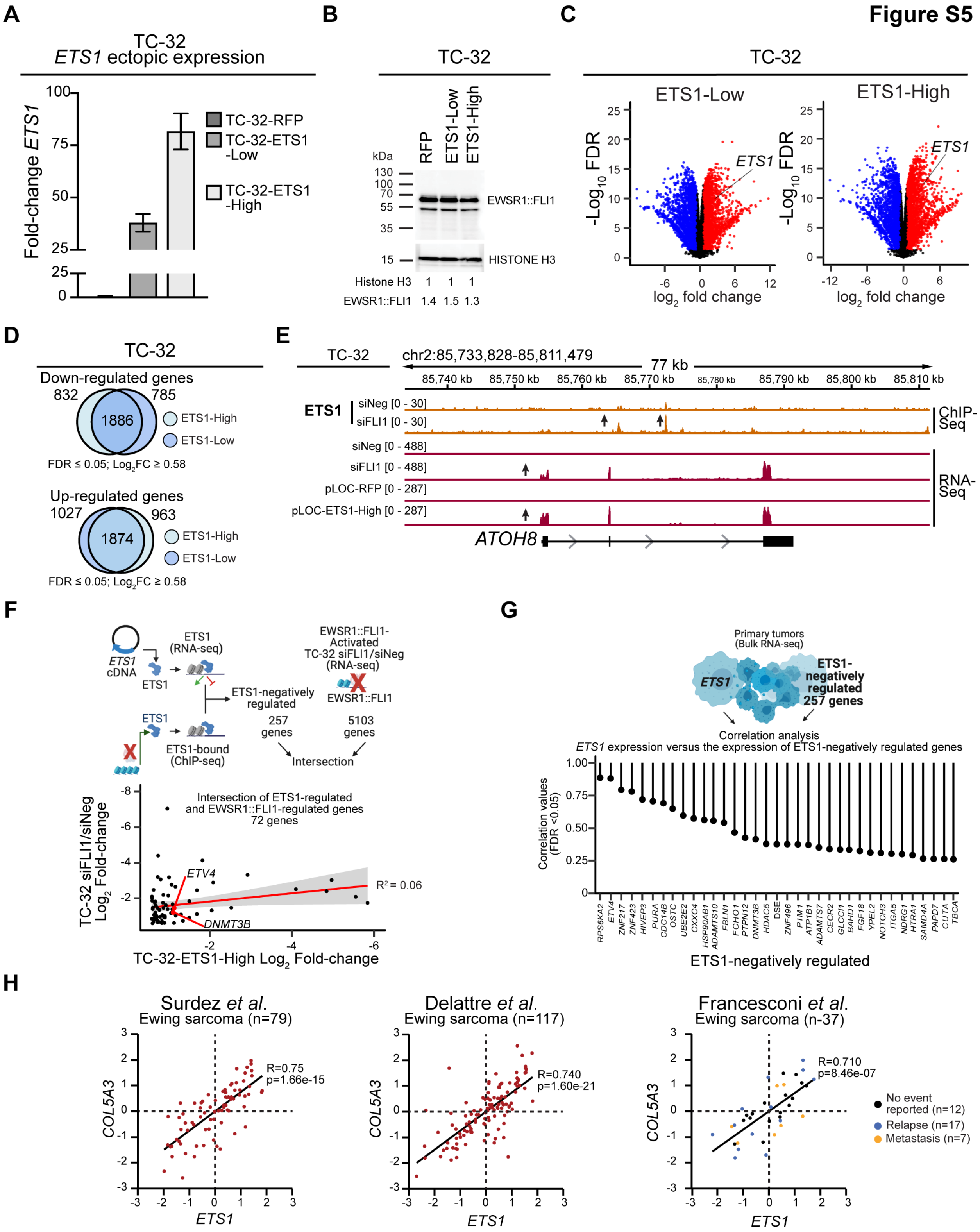

**Figure S5: Ectopic expression of ETS1 has transcriptome-wide effects.** (**A**) qRT-PCR analysis of *ETS1* expression exhibited by the control and the two single-cell clones expressing an *ETS1* cDNA (NM_01143820; isoform 1). Results are shown as mean ± SEM of three replicates. (**B**) Immunoblots of whole-cell lysates prepared from the indicated cell lines were analyzed using antibodies against the indicated proteins. (**C**) Changes in gene expression were observed following the ectopic expression of *ETS1*. (**D**) The co-occurrence of genes that exhibit significant changes in expression following the ectopic expression of *ETS1* in the indicated TC-32 single-cell clones (see also **Supplemental Table S8**). (**E**) The *ATOH8* locus of TC-32 cells showing ETS1 ChIP-seq peaks (siNeg or siFLI1-transfected) and RNA-seq read depth (siNeg or siFLI1-transfected cells or modified – RFP or ETS1-High). **(F)** A model depicting the analysis of the expression of ETS1-negatively regulated genes and genes that exhibit a decrease in expression following the depletion of EWSR1::FLI1 and the result of this analysis is presented as a scatter plot. **(G)** A model depicting the correlation analysis of *ETS1* expression and the expression of ETS1-negatively regulated genes in EWS tumors and the results of this analysis using the gene expression profiles of 79 EWS tumors (19) focused on significantly correlated genes (FDR <0.05). (**H**) The correlation of *ETS1* and *COL5A3* mRNA levels in EWS tumors reported previously in the indicated studies: GSE142162 79 samples (24); GSE34620 (117 tumors) (22); GSE12102 (37 tumors) (23), analyzed and plotted using the R2 Genomics Visualization platform (20).

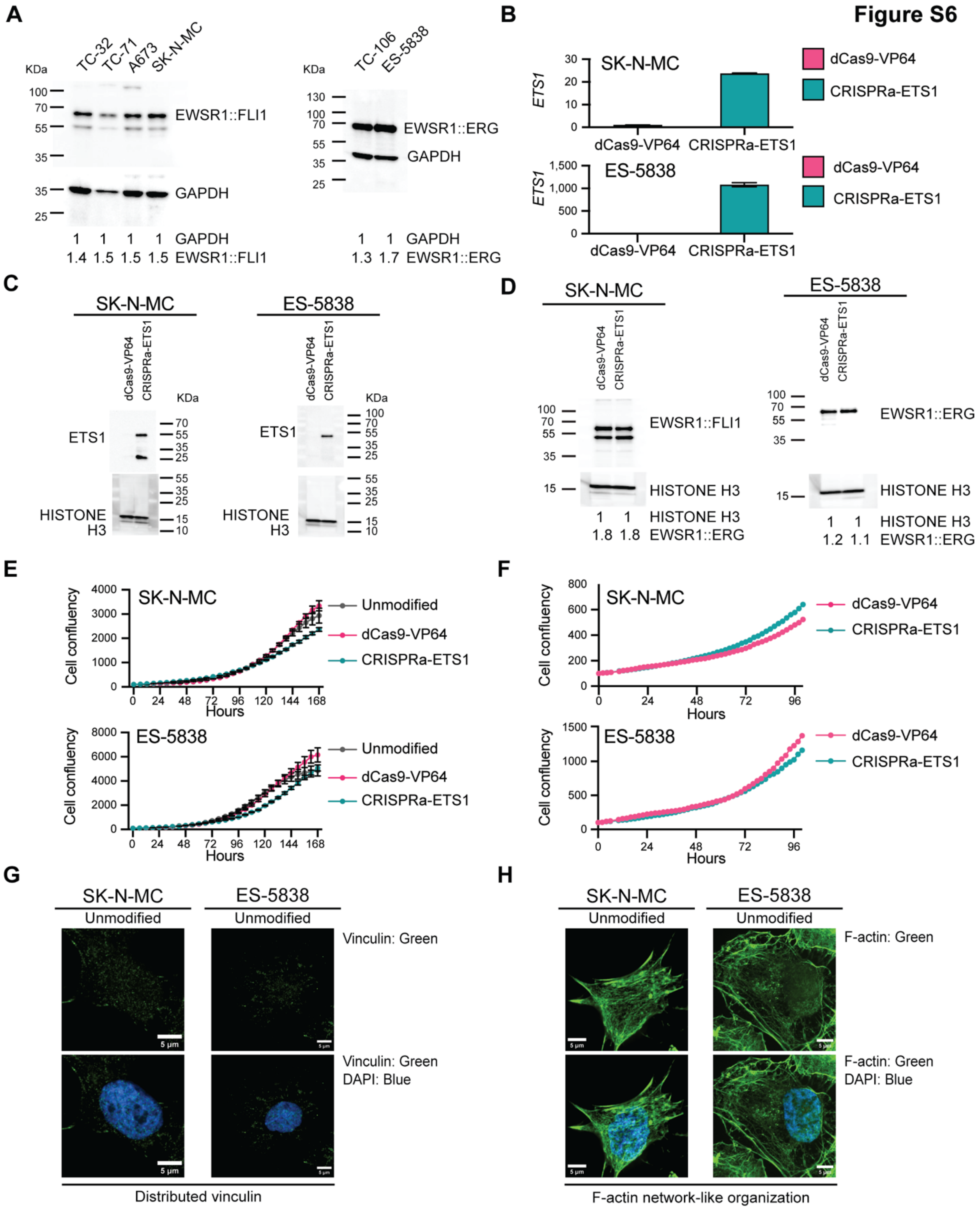

**Figure S6: ETS1 promotes the migration of EWS cells.** (**A**) Immunoblots of whole-cell lysates prepared from the indicated cell lines and analyzed using antibodies against the indicated proteins. (**B**) The expression of *ETS1* assessed by qRT-PCR analysis in the indicated unmodified and modified EWS cell lines. Results are shown as mean ± SEM of three replicates, except for duplicate ES-5838-dCas9-VP64 samples. (**C**) Immunoblot analysis of whole-cell lysates prepared from the indicated unmodified and modified EWS cell lines and analyzed using the antibodies against the indicated proteins. (**D**) Immunoblot analysis of whole-cell lysates prepared from the indicated unmodified and modified EWS cell lines and analyzed using the antibodies against the indicated proteins. (**E**) Cell confluency (area of confluency normalized to 0 hrs) over time of the indicated unmodified and modified (dCas9-VP64 and CRISPRa-ETS1) EWS cell lines. Each data point indicates the mean ± SEM of four replicates representing three independent experiments. (**F**) Cell confluency (area of confluency normalized to 0 hrs) over time of dCas9-VP64 and CRISPRa-ETS1 EWS cell lines employed in the analysis of cell movement (**Fig. 6B, 6C** and **videos 1** - **4**). The data shown indicates the mean ± SEM of 46 replicates. (**G)** Super-resolution merged images of unmodified SK-N-MC and ES-5838 cells analyzed using an antibody against vinculin (green) and the DAPI stain (blue). (**H**) Super-resolution merged images of unmodified SK-N-MC and ES-5838 cells analyzed using an antibody against F-actin (green) and the DAPI stain (blue).

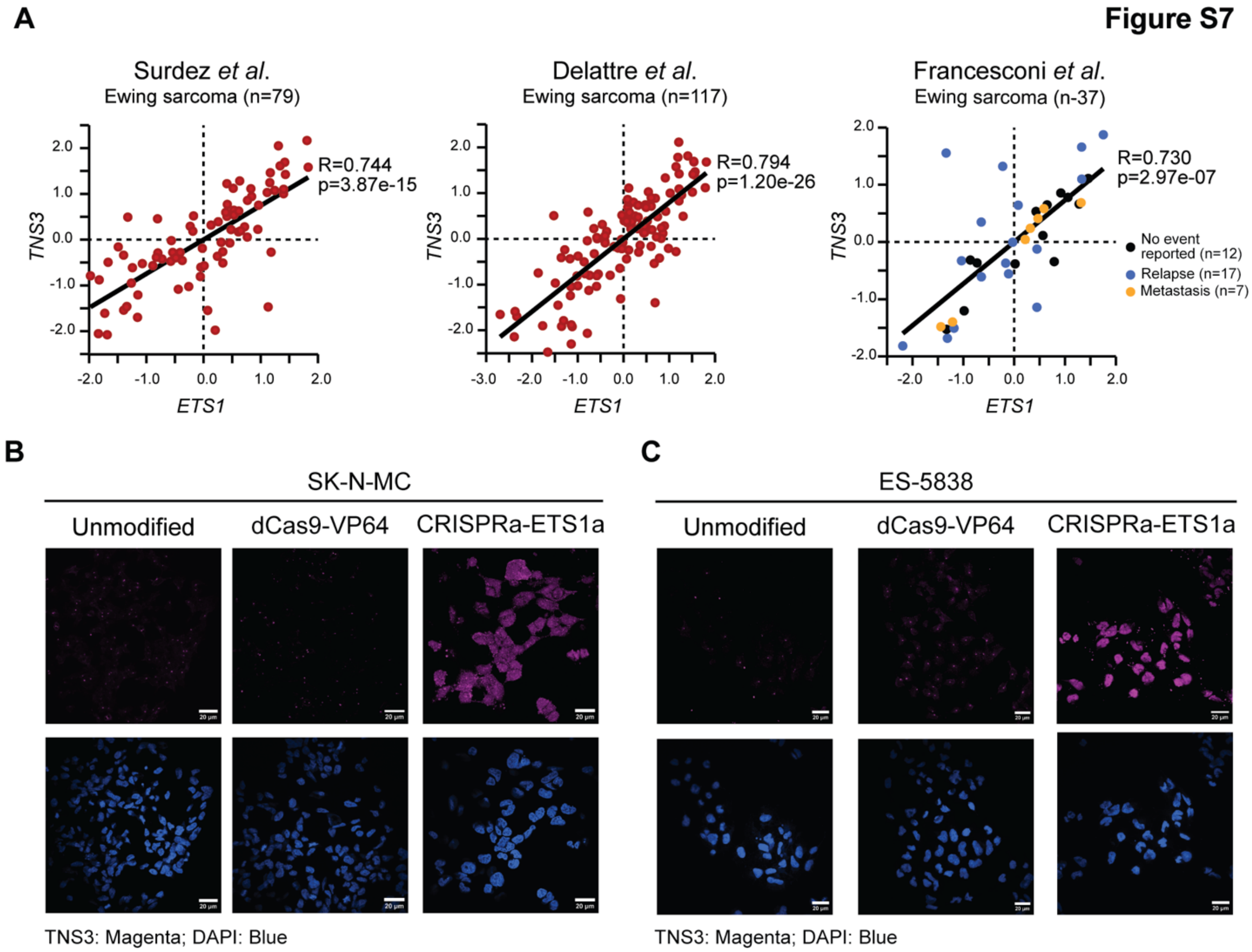

**Figure S7: ETS1 regulates the expression of TENSIN3.** (**A**) The correlation of *ETS1* and *TNS3* mRNA levels in EWS tumors reported previously in the indicated studies: GSE142162 79 samples (24); GSE34620 (117 tumors) (22); GSE12102 (37 tumors) (23), analyzed and plotted using the R2 Genomics Visualization platform (20). (**B**) Confocal single-channel images of DAPI (blue) and TNS3 (magenta) in unmodified and modified SK-N-MC cells corresponding to the merged images shown in **Fig. 7G**. (**C**) Confocal single-channel images of DAPI (blue) and TNS3 (magenta) in unmodified and modified ES-5838 cells corresponding to the merged images shown in **Fig. 7G**.

**Figure S8**

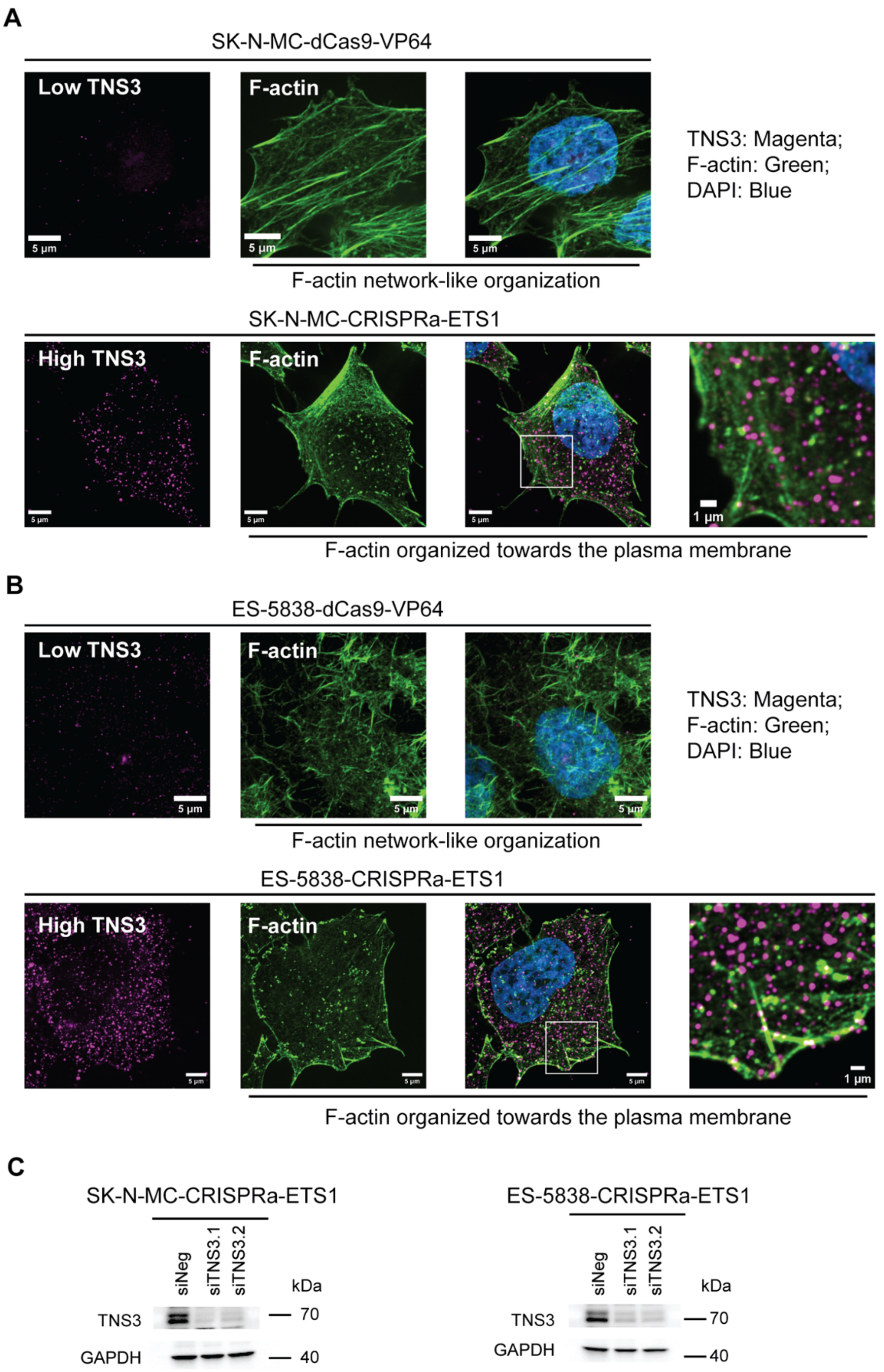

**Figure S8: EWS cells expressing ETS1 exhibit TENSIN3-dependent changes in cytoskeletal organization.** (**A**) Super-resolution single-channel and merged images (including a zoomed inset) of the indicated modified SK-N-MC cells analyzed using antibodies against TNS3 (magenta), F-Actin (green), and the DAPI stain (blue). (**B**) Super-resolution single-channel and merged images (including a zoomed inset)

of the indicated modified ES-5838 cells analyzed using antibodies against TNS3 (magenta), F-Actin (green), and the DAPI stain (blue). (**A, B**) Cell images, scale bar = 5 μm, zoomed inset, scale bar = 1 μm; images represent 30 images across two independent replicates. (**C**) Immunoblots of whole-cell lysates prepared from cells transfected with the indicated siRNAs for 48 hrs and analyzed using the antibodies against the indicated proteins. Studies examining TNS3 function in mediating changes in the cytoskeletal organization used the siRNA designated siTNS3.2 (**Fig. 8C-F**).

### Supplemental Materials and Methods

**RNA sequencing:** For paired-end RNA sequencing (RNA-seq), total RNA was extracted (Maxwell 16 LEV simplyRNA purification kit, Promega, Madison, WI), and poly(A) selected RNA and sequencing libraries prepared using a TruSeq Standed mRNA LT Sample Preparation Kit following the manufacturer's instructions (Illumina, San Diego, CA). Libraries were sequenced using a Nextseq 2000 instrument (Illumina). The CCR Collaborative Bioinformatics Resource (CCBR) RNA-seq pipeline was used for RNA-seq analysis (https://bioinformatics.ccr.cancer.gov/ccbr/pipelines-software/ccbr-pipeliner/). FeatureCounts in the Subread v.2.0.3 package (1) was used to summarize gene level reads by counting reads that overlapped the GRCh38/hg38 annotated gene exons. Limma v.3.38.3 was employed for gene count normalization of each sample using RSEM (RNA-Seq by Expectation-Maximization) $\log_2$ transcript per million (TPM) values and the quantification of differential gene expression comparing control and experimental conditions (2,3). Metascape (https://metascape.org) was used for the gene ontology (GO) analysis of upregulated and downregulated genes in three Ewing sarcoma cell lines (TC-32, TC-71, and A673) following the depletion of EWSR1::FLI1. For the summary of GO-terms related to biological processes shown in **Figure S1C**, we focused on those terms that appeared in both Metascape and GSEA database, plotting the P values from the Metascape analysis.

#### Chromatin Immunoprecipitation (ChIP) and CUT&RUN analysis

**ChIP-based analysis:** ChIP assays were performed using SimpleChIP plus Enzymatic Chromatin IP kit (9005S; Cell Signaling Technology, Danvers, MA) following the manufacturer's protocol. Nuclei were ruptured by sonication (Branson SFX250 Digital Sonifier; Branson Ultrasonics, Brookfield, CT) using the following settings, 30% energy, 30 sec ON, 30 sec OFF, total ON time 3 minutes (mins), and insoluble fractions were removed by centrifugation. ChIP inputs (1%) from clarified sheared chromatin were de-crosslinked by adding elution buffer supplemented with 6 µl of NaCl (5 M) and 2 µl Proteinase K (20mg/ml), following incubation for 16 hours (hrs) at 65°C. The remaining clarified sheared chromatin was incubated with primary antibodies (5 µg of ETS1 or 3 µg of H3K27ac and H3K4me3) overnight at 4 °C. Drosophila chromatin and an antibody against a Drosophila specific histone variant H2Av (Active Motif, Carlsbad, CA) were used as calibration controls per the manufacturer's recommendations. Following incubation for 12-18 hrs, the ChIP pulldowns were performed using protein G magnetic beads and eluted in 150 µl elution buffer supplemented with 6 µl NaCl (5M) and 2 µl Proteinase K (20 mg/ml) and de-crosslinked for 16 hrs at 65°C. The enrichment for binding of specific regions was assessed relative to the input DNA. ChIP-seq library preparations and sequencing were performed using either Swift Accel-NGS® 2S Plus DNA Library Kit (21096, Swift Biosciences) or TAKARA SMARTer ThruPLEX DNA-Seq Library Prep Kit (R400676, Takara Bio) following manufacturer's protocols. Two independent replicates of each ChIP samples were prepared except H3K27ac which was prepared in triplicate.

**CUT&RUN analysis:** Cleavage Under Target and Release Using Nuclease (CUT&RUN) was carried out as previously described (4) with slight modifications. In brief, 5 x $10^5$ cells/condition were harvested, washed, and bound to activated Concanavalin A (ConA) conjugated paramagnetic beads (21-1401; EpiCypher, Durham, NC). The ConA beads–cell complex was resuspended in antibody buffer (20 mM HEPES, 150 mM NaCl, 0.5 mM Spermidine, protease inhibitor, 0.01% digitonin, and 2 mM EDTA), and incubated with the appropriate antibody (**Supplemental Table S1**) overnight at 4°C. Following incubation, pAG-MNase (15-1016; EpiCypher) was added to each CUT&RUN reaction to allow binding to the antibody-labelled chromatin. Targeted chromatin was digested and released by the addition of CaCl2 (supplemented with S. cerevisiae spike-in, 29987L; Cell Signal Technology). Fragmented chromatin was purified using CUTANA Purification kit (14-0050; EpiCypher) and purified DNAs quantified using a Qubit fluorometer (Thermo Fisher Scientific). The CUTANA CUT&RUN Library Prep kit (14-1001, EpiCypher) was used for library preparation following manufacturer's instructions. Briefly, end repair 5' phosphorylation and 3' dA-Tailing was performed by adding end repair master mix (End Prep Buffer and End Prep Enzyme) to the purified DNA. The thermocycler conditions were set to 20°C for 20 min and 65°C for 30 min to prevent thermal degradation of the shortest fragments. Adapter ligation and Uracil-specific excision was performed at 4°C by adding the ligation master mix (ligation mix and ligation enhancer) and U-excision enzyme. A post-ligation DNA cleanup was performed using SPRIselect beads (Beckman Coulter, Brea, CA) at a ratio of 1:1 sample volume. Adapter-ligated DNA was amplified using the High Fidelity 2X PCR master (New England Biolabs) and indexed using selected i5 and i7 indexing primers (14-1001, EpiCypher) with the following thermocycler conditions; 98°C for 45 sec, 14 cycles of 98°C for 15 sec, 60°C for 10 sec and final extension at 72°C for 60 sec. Post-PCR clean-up was performed on amplified libraries with SPR1select beads 0.9X left size selection then washed twice gently in 80% ethanol and eluted in 0.1X TE buffer. Sequencing was performed using standard protocols.

**ChIP and CUT&RUN bioinformatic analysis:** For analysis of ChIP-seq data, reads were trimmed for adapters using CutAdapt (5). Trimmed reads were aligned to the GRCh38/hg38 human genome and the *Drosophila melanogaster* dm6 genomes using BWA mem v.0.7.17 (6).

PCR duplicates were removed using Picard v.2.17.11 SamToFastq (for blacklist read removal) and MarkDuplicates (to remove PCR duplicates) (https://broadinstitute.github.io/picard/). Peak calling was performed against input control using model-based MACS2 v2.1.1 (7), with the cut-off q-value < 0.01. Narrow peaks were identified for all ChIP-seq samples. Peaks were annotated with the closest hg38 genes using Uropa v.4.0.2 (8), and motif enrichment analysis was performed using Homer v.4.10.1 (9). DeepTools v3.5.1 suite (11) was used to created bigwig files and the bamcoverage function was used to create reads per genomic content (RPGC) or reads per million (RPM)-normalized data to account for differences in sequencing depth. For analysis of CUT&RUN data, the CCBR ChIP-seq pipeline (https://github.com/CCBR/CARLISLE) was used. MACS2 narrow was used to call peaks for EWSR1::FLI1 and H3K27ac, while MACS2 broad was used to call peaks for H3K9me3 and H3K27me3 using CUTANA™ Rabbit IgG as a control. CUT&RUN samples were normalized to *Saccharomyces cerevisiae sacCer3* spike-in control. Gene promoter regions were defined as – 3 kb and + 1 kb intervals around the hg38 gene transcription start site (TSS). Distal regions were defined outside the promoter regions.BEDTools v.2.30.0 (10) was used for genomic region analyses (merging and intersection) and a combination of bedtools and functions from the deepTools v.3.5.1 suite (computeMatrix, plotHeatmap, and plotProfile (11) were used to plot heatmaps and summary profiles. The Integrative Genomic Viewer (IGV) v2.15.1 (12) platform was used to visualize gene-specific datasets.

**Immunoblotting and Immunofluorescence.** For immunoblotting, membranes were developed using SuperSignal™ West Pico PLUS Chemiluminescent Substrate (ThermoFisher Scientific) and visualized using an Omega Lum™ C Imaging System (Aplegen). Quantification of relative protein expression was performed in ImageJ. In brief, a region of interest (ROI) was defined around the largest signal and the mean gray value measured. Maintaining the ROI constant, intensity values were obtained for all signals in each lane including the loading controls and ten background values in grey scale. Values were exported to Microsoft Excel and the intensity values for each signal were first background subtracted and further normalized to the loading control and depicted as shown. For analysis using IF, cells grown on round coverslips for 72 – 96 hrs were washed once with PBS and fixed with 4% paraformaldehyde for 3 mins. Cells were washed thrice with PBS and permeabilized for 5 mins with a 0.5% Triton X-100 solution (Sigma Chemical, St. Louis, MO) in PBS. Cells were washed thrice with PBS and blocked in 10% normal goat serum for an hour (h) before incubating with either primary or conjugated antibodies overnight at the dilutions detailed in **Supplemental Table S1**, and DAPI to define nuclei.

**Supplementary Table S1: Reagents and resources**

| REAGENT or RESOURCE | SOURCE | IDENTIFIER |
|---|---|---|
| **Antibodies** | | |
| Rabbit anti-FLI1*<br>Immunoblot – 1:1000 dilution | Abcam | ab15289 |
| Rabbit anti-FLI1*<br>CUT&RUN – 1:50 dilution | Abcam | ab15289 |
| Rabbit anti-ERG<br>(Immunoblot 1:1000) | Abcam | ab92513 |
| Rabbit anti-ETS1<br>Immunoblot – 1:1000 dilution | Cell Signaling Technology | 14069S |
| Rabbit anti-ETS1<br>ChIP-seq – 5 µg | Cell Signaling Technology | 14069S |
| Rabbit anti-SNAI2 (SLUG)<br>Immunoblot – 1:300 dilution | Cell Signaling Technology | 9585S |
| Rabbit anti-JUNB<br>Immunoblot – 1:1000 dilution | Cell Signaling Technology | 3753SS |
| Rabbit anti-RUNX2<br>Immunoblot – 1:2000 dilution | Cell Signaling Technology | 12556S |
| Rabbit anti-alpha TUBULIN<br>Immunoblot – 1:5000 dilution | Cell Signaling Technology | 2144S |
| Rabbit anti-H3K27ac<br>ChIP-seq – 3 µg | Abcam | ab4729 |
| Rabbit anti-H3K27ac<br>CUT&RUN – 1:50 dilution | Abcam | ab4729 |
| Rabbit anti-H3K27ac<br>ChIP-seq – 3 µg | Active Motif | 39133 |
| Rabbit anti-H3K27me3<br>CUT&RUN – 1:50 dilution | Cell Signaling Technology | 9733S |
| Rabbit anti-H3K9me3<br>CUT&RUN – 1:50 dilution | Abcam | ab8898 |
| Rabbit anti-H3K4me3<br>ChIP-seq – 3 µg | EpiCypher | 13-0041 |
| Rabbit IgG Isotype Control<br>CUT&RUN – 1:25 dilution | Cell Signaling Technology | 66362S |
| Normal Rabbit IgG<br>ChIP-qPCR – 2 µg | Cell Signaling Technology | 2729S |
| Rabbit anti-TENSIN 3<br>Immunoblot – 1:1000 | ThermoFisher | PA5-63112 |
| Rabbit anti-TENSIN 3<br>IF – 1:500 | ThermoFisher | PA5-63112 |
| Rabbit anti-GAPDH<br>Immunoblot – 1:3000 | Santa Cruz | sc-47778 |
| Goat anti-rabbit – 1:3000 to 1:5000 | Cell Signaling Technology | 7074S |
| Goat anti-mouse – 1:3000 to 1:5000 | Cell Signaling Technology | 7076S |
| Alexa Fluor™ 488 Phalloidin (F-actin)<br>IF- 1:1000 dilution | Thermo Fisher Scientific | A12379 |

| | | |
|---|---|---|
| Vinculin (monoclonal) AlexaFluor 488 conjugated<br>IF-1:1000 dilution | Thermo Fisher Scientific | 53-9777-80 |
| Anti-mouse IgG (H+L), F(ab')$_2$ Fragment (Alexa Fluor® 647 Conjugate<br>IF-1:1000 dilution | Cell Signaling Technology | 4410 |
| Anti-rabbit IgG (H+L), F(ab')2 Fragment (Alexa Fluor® Conjugate<br>IF-1:1000 dilution | Cell Signaling Technology | 4413 |
| *The use of an anti-FLI1 antibody is an accepted approach to identify EWSR1::FLI1 binding (Riggi *et al.* Cancer Cell 2014;26:668-81; Adane *et al.* Cancer Cell 2021;39:827-44 e10; Showpnil *et al.*. Nucleic Acids Res 2022; 50:9814-37; Gao *et al.* Nat Cell Biol 2023;25:298-308**;** Lu *et al.* Nat Cell Biol 2023;25:285-97. | | |
| **Cell lines** | | |
| A673 (DMEM) | ATCC | CRL-1598™<br>STR October 2019 |
| TC-32 (RPMI 1640) | Pediatric Oncology Branch, CCR | STR October 2019 |
| ES-5838 (RPMI 1640) | Pediatric Oncology Branch, CCR | STR November 2019 |
| ES-5838-dCas9-VP64 | This study | STR November 2023 |
| ES-5838-CRISPRa-ETS1 | This study | STR November 2023 |
| TC-106 (RPMI1640) | Pediatric Oncology Branch, CCR | STR October 2019 |
| SK-N-MC (RPMI 1640) | Javed Khan, Genetics Branch, CCR | STR October 2019 |
| SK-N-MC-dCas9-VP64 | This study | STR November 2023 |
| SK-N-MC- CRISPRa-ETS1 | This study | STR November 2023 |
| TC-71 (RPMI 1640) | Pediatric Oncology Branch, CCR | STR October 2019 |
| HEK-293T (DMEM) | ATCC | STR November 2019 |
| Chemicals and commercial assays | | |
| BCA Protein assay kit | Thermo Fisher Scientific | J63283.QA |
| CUTANA™ ChIP/CUT&RUN Kit | EpiCypher | 14-1048 |
| DAPI (4′,6-diamidino-2-phenylindole) 1mg/ml solution | ThermoFisher Scientific | 62248 |
| Trans-Lentiviral ORF Packaging Kit with Calcium Phosphate Transfection Reagent | Dharmacon/Horizon Discovery | TLP5912 |
| DMEM, high glucose, pyruvate | Thermo Fisher Scientific | 11995073 |
| Fetal Bovine Serum (FBS), qualified, heat inactivated, United States | Thermo Fisher Scientific | 16140071 |

| | | |
|---|---|---|
| Halt™ Protease and Phosphatase Inhibitor Cocktail, EDTA-free (100X) | Thermo Fisher Scientific | 78445 |
| Lipofectamine™ 3000 Transfection Reagent | Thermo Fisher Scientific | L3000015 |
| Lipofectamine™ RNAiMax Transfection Reagent | Thermo Fisher Scientific | 13778150 |
| Maxwell Genomic DNA extraction kit | Promega | AS1140 |
| Maxwell RNA extraction kit | Promega | AS1390 |
| MycoAlert Mycoplasma Detection Kit | Lonza | LT07- 218 |
| Normal Goat Serum | Cell Signaling Technology | 5425S |
| Normal Goat Serum (10%) | ThermoFisher | 50062Z |
| Nucleofector Kit R | Lonza | VCA-1001 |
| PageRuler™ Plus Pre-stained Protein Ladder, 10 to 250 kDa | Thermo Fisher Scientific | 26619 |
| PBS | Thermo Fisher Scientific | 70011044 |
| PEG Virus Precipitation Kit | Abcam | Ab102538 |
| pMD2.G | Addgene | 12259 |
| psPAX2 | Addgene | 12260 |
| Plasmocin Prophylactic | InvivoGen | ant-mpp |
| RIPA Buffer | Cell Signaling Technology | 9806 |
| RPMI 1640 Medium | Thermo Fisher Scientific | 11875119 |
| SuperSignal™ West Pico PLUS Chemiluminescent Substrate | Thermo Fisher Scientific | 34580 |
| SYBR™ Safe DNA Gel Stain | Thermo Fisher Scientific | S33102 |
| SimpleChIP® Plus Enzymatic Chromatin IP Kit | Cell Signaling Technology | 9005S |
| Puromycin Dihydrochloride (10mg/mL) | Thermo Fisher Scientific/Gibco | A1113803 |
| Blasticidin S HCl (10 mg/mL) | Thermo Fisher Scientific/Gibco | A1113903 |

| **PCR Primers** | | |
|---|---|---|
| *EWSR1::FLI1 forward,* ACCCCAAACTGGATCCTACAG | Thermo Fisher Scientific | NA |
| *EWSR1::FLI1 reverse,* GGCCGTTGCTCTGTATTCTTAC | Thermo Fisher Scientific | NA |
| *EWSR1::ERG forward,* CTTCCACAGTGCCCAAAACT | Thermo Fisher Scientific | NA |
| *EWSR1::ERG reverse,* AACTGCCAAAGCTGGATCTG | Thermo Fisher Scientific | NA |
| Hs_BHLHE40_1 | Qiagen | QT00035210 |
| *EPAS1 forward,* TCCAACAAGCTGAAGCTGAA | Thermo Fisher Scientific | NA |
| *EPAS1 reverse,* TCATCCGTTTCCACATCAAA | Thermo Fisher Scientific | NA |
| *ETS1 forward,* AAGACTGCTTTCTCGAGCTG | Thermo Fisher Scientific | NA |
| *ETS1 reverse,* TAGGCTGGGTTGACTCCATT | Thermo Fisher Scientific | NA |
| Hs_HEYL_1 | Qiagen | QT00073052 |
| Hs_JUN_1 | Qiagen | QT00242956 |

| | | |
|---|---|---|
| *JUNB forward,* TGGAACAGCCCTTCTACCAC | Thermo Fisher Scientific | NA |
| *JUNB reverse,* AGGCTCGGTTTCAGGAGTTT | Thermo Fisher Scientific | NA |
| Hs_PPARD_1 | Qiagen | |
| *RUNX2 forward*, CGGAATGCCTCTGCTGTTAT | Thermo Fisher Scientific | NA |
| *RUNX2 reverse*, TGAAGACGGTTATGGTCAAGG | Thermo Fisher Scientific | NA |
| *SNAI2 forward*, CGAACTGGACACACATACAGTG | Thermo Fisher Scientific | NA |
| *SNAI2 reverse*, CTGAGGATCTCTGGTTGTGGT | Thermo Fisher Scientific | NA |
| *ETS1_A (EWSR1::FLI1 ChIP) forward 1,* GGGCAGACAGGAACCAGAA | Thermo Fisher Scientific | NA |
| *ETS1_A (EWSR1::FLI1 ChIP) reverse 1,* CTGGAGTGGTGTGTAAGGCT | Thermo Fisher Scientific | NA |
| *ETS1_A (EWSR1::FLI1 ChIP) forward 2,* GCACAGCTGTTTAAAGACGC | Thermo Fisher Scientific | NA |
| *ETS1_A (EWSR1::FLI1 ChIP) reverse 2,* TCTTTGCCACAGATGTCCCT | Thermo Fisher Scientific | NA |
| *ETS1_B (EWSR1::FLI1 ChIP) forward 1,* TCCAATGAGCCCTTGACCAT | Thermo Fisher Scientific | NA |
| *ETS1_B (EWSR1::FLI1 ChIP) reverse 1,* GCAATCCCTTTTGTCTGGCA | Thermo Fisher Scientific | NA |
| *ETS1_B (EWSR1::FLI1 ChIP) forward 2,* AGCCAGGCCTGAGAAAGAAG | Thermo Fisher Scientific | NA |
| *ETS1_B (EWSR1::FLI1 ChIP) reverse 2,* GTCAAGGGCTCATTGGAACC | Thermo Fisher Scientific | NA |
| *ETS1_C (EWSR1::FLI1 ChIP) forward,* GCTGACAGGAAGGCTCTATT | Thermo Fisher Scientific | NA |
| *ETS1_C (EWSR1::FLI1 ChIP) reverse,* TGGGAATGAGAAGGGCAGAA | Thermo Fisher Scientific | NA |
| *ETS1_D (EWSR1::FLI1 ChIP) forward,* GAGGAGCAGTGCGTGGAG | Thermo Fisher Scientific | NA |
| *ETS1_D (EWSR1::FLI1 ChIP) reverse,* GACCAAGCCCTCAAGAATGC | Thermo Fisher Scientific | NA |
| *COL5A3 forward (ETS1 ChIP)* GGGGCTCCTGAGAAATTCCT | Thermo Fisher Scientific | NA |
| *COL5A3 reverse,* GCAGGAGTGGAAGTTGCTC | Thermo Fisher Scientific | NA |
| *TNS3 I forward,* GCTGAGCCGCCAAACACT | Thermo Fisher Scientific | NA |
| *TNS3 I reverse,* CGGCGCGTCGGTAATTCC | Thermo Fisher Scientific | NA |
| *TNS3 II forward,* TGCCCGTCAGAAGTCAGAC | Thermo Fisher Scientific | NA |
| *TNS3 II reverse,* TTCGTGTCAGATGGAGGGAC | Thermo Fisher Scientific | NA |

| **RNAi** | | |
|---|---|---|
| Negative control (siNeg) | Qiagen | SI03650318 |
| FLI1-fusion (siFLI1)<br>5'-CAAACGAUCAGUAAGAAUAtt-3' | ThermoFisher Scientific<br>Silencer Select | S5266 |
| ERG-fusion (siERG)<br>5'-CCACAGUGCCCAAAACUGAtt-3' | ThermoFisher Scientific<br>Silencer Select | s4812 |
| TNS3 siRNA 1 (siTNS3.1)<br>5'-GCUCAUUCAUUGUUCGAGAtt-3' | ThermoFisher Scientific<br>Silencer Select | s34875 |
| TNS3 siRNA 2 (siTNS3.2)<br>5'-CUUACGAAGCUUAACCCAAtt-3' | ThermoFisher Scientific<br>Silencer Select Silencer Select | s34876 |

| **CRISPRa Plasmids** | | |
|---|---|---|
| Lenti dCas9-VP64_Blast | Addgene, Watertown, MA | 61425 |
| *ETS1-Calabrese-SetA-2,* (sgRNA)<br>5'-CCTCTTTAGGGCGTTTCTCG-3' | Addgene, Watertown, MA | NA |
| *ETS1-Calabrese-SetB-1,* (sgRNA)<br>5'-CCGCGAGAAACGCCCTAAAG-3' | Addgene, Watertown, MA | NA |
| *ETS1-Calabrese-SetB-2,* (sgRNA)<br>5'-CGAGAAACGCCCTAAAGAGG-3' | Addgene, Watertown, MA | NA |

| Software and Algorithms | | |
|---|---|---|
| Prism 9 | GraphPad Software | https://www.graphpad.com/scientific-software/prism/ |
| Benchling | Benchling Inc | Volume 1, 2023 |
| RStudio | | RStudio Version 1.4.1717 © 2009-2021 RStudio, PBC |
| Geneious Prime with minimap2 plugin | BioMatters Ltd | Geneious Prime® 2023.0.1 Build 2022-11-28 12:49 |
| Fiji | | V1.53t |
| NIS software | Nikon | V5.3 |

| Ewing sarcoma cell lines included in the cited RNA-seq dataset |
|---|
| 6647 |
| A673 |
| CHLA10 |
| CHLA25 |
| TTC466 |
| CHLA258 |
| CHLA32 |
| TTC475 |
| CHLA352 |
| COGE352 |
| CHLA9 |
| CHP100L |
| ES1 |
| ES2 |
| ES3 |

| |
|---|
| ES4 |
| ES6 |
| ES7 |
| ES8 |
| EW8 |
| NCIEWS5000 |
| RDES |
| SKES1 |
| SKNEP1 |
| SKNMC |
| SKPNETLI |
| TC106 |
| TC138 |
| SKNLO |
| TC167 |
| TC177 |
| TC215 |
| TC233 |
| TC240 |
| TC244 |
| TC248 |
| TC253 |
| TC32 |
| TC487 |
| TTC547 |
| TC4C |
| TC71 |
